## Supplementary material for "Gene expression correlates of social evolution in coral reef butterflyfishes"

Department of Biology

Stanford University

371 Jane Stanford Way

Stanford, CA 94305

Journal: Proceedings of the Royal Society, B

DOI: 10.1098/rspb.2020.0239

**Supplementary Methods S1**: Detailed qPCR gene expression analysis

Frozen brains were coronally sectioned on a cryostat at 100µm, thaw mounted onto Superfrost Plus slides (Fisher Scientific) and stored at -80˚C. Brain regions were identified using a butterflyfish brain atlas (Bauchot, 1989; Dewan and Tricas, 2014) and manually extracted at -30˚C using a hand-held micro-punching device (50mm diameter; Stoelting, model # 57401) (**Fig. 2 inset**). Punches from individual brain regions were pooled in RNA*later*™ (ThermoFischer, cat no. AM7020) at 4˚C overnight, and then stored at -20˚C for up to one week. Tissue punches were transferred into a lysis buffer and homogenized by passing through a 21-gauge needle 15 times. RNA was extracted using an E.Z.N.A® HP Total RNA kit (Omega Bio Tek, cat no # R6812-02) according to the manufacturer's instructions, and stored at -80˚C prior to cDNA synthesis. RNA was reverse transcribed into cDNA using Superscript III reverse transcriptase (Life Technologies) and gene-specific primers (see Table S1 for primer sequences). Residual primers and salts from reverse transcription were removed using an E.Z.N.A ® Tissue DNA purification kit (Omega, product # D3396-02), and cDNA was stored at -20˚C for up to 10 days prior to qPCR.

The whole brain transcriptome of *C. lunulatus* was sequenced as a reference for designing species specific cloning primers for target genes. One *C. lunulatus* brain was taken out of RNAlater, rinsed in 1X phosphate buffered saline (PBS) and placed immediately in Trizol (Life Technologies, Grand Island, NY) where RNA was extracted according to manufacturer instructions. Poly-adenylated RNA was isolated from each sample using the NEXTflex PolyA Bead kit (Bioo Scientific, Austin, TX, USA). Lack of contaminating ribosomal RNA was confirmed using the Agilent 2100 Bioanalyzer. A strand specific library was prepared using the dUTP NEXTflex RNAseq kit (Bioo Scientific), which includes a magnetic bead-based size selection of roughly 350 bp. The library was pooled in equimolar amounts with samples from an unrelated study after library quantification using both quantitative PCR with the KAPA Library Quantification Kit (KAPA Biosystems, Wilmington, MA, USA) and the fluorometric Qubit dsDNA high sensitivity assay kit (Life Technologies), both according to manufacturer instructions. Libraries were sequenced on an Illumina HiSeq 2000 to obtain paired-end 100bp reads. We first corrected errors in the Illumina reads using Rcorrector (parameters: run_rcorrector.pl -k 31) and then applied quality and adaptor trimming using Trim Galore! (http://www.bioinformatics.babraham.ac.uk/projects/trim_galore/; parameters: trim_galore --paired --phred33 --length 36 -q 5 --stringency 5 --illumina -e 0.1). After filtering and trimming, a total of 64,795,096 paired reads remained for de novo assembly. We created a *C. lunulatus de novo* transcriptome assembly using Trinity (parameters: --seqType fq --SS_lib_type RF). The raw Trinity assembly produced 376,338 contigs (N50: 1148 bp). Raw data and the *C. lunulatus* transcriptome is available on read archive (submission pending acceptance).

Using the *C. lunulatus* transcriptome, primers were designed to clone target gene sequences. Sequences of *OTR*, *V1aR*, *D1R*, *D2R* and *MOR* genes were then used to design qPCR primers that flanked exon boundaries using *Danio rerio* and *Stegastes partitus* genomes as a reference (see Table S1). Prior to qPCR, primer sets and instrument cycling parameters were empirically optimized on standard curves using several metrics of quality control (i.e., assay amplification $R^2$ value of at least 0.95, assay slope of approximately -3.3, assay melting curve that only produced a single amplicon peak, no amplicon signal in the no template control (NTC) or no reverse-transcriptase control (NRTC)). Quantitative PCR was then performed on each sample using a reaction mixture and qPCR cycling instrument (CFX380) recommended by the enzyme manufacturer (see Table S2 for parameters). Samples were run in technical triplicate on 384 well qPCR plates with standard curves in order to determine assay efficiency. Not all regions of each brain were measured for gene expression due to insufficient tissue available.

**Supplementary Methods S2**: R script for analysis of qPCR data

*Chaetodon lunulatus*: *Social system X sex:*

```
# 1. install.packages("MCMC.qpcr")

library(MCMC.qpcr)

# 2. read in data

data <- read.csv(file="Nowicki_C_Lun_M&F_data_ESM.csv")

data$count <- as.numeric(data$count) # force counts to be numeric

head(data) # allows you to view your data to ensure it was imported correctly

# 3. sub-set data into different brain regions and store them in separate data frames, in order to analyse each brain region separately

levels(data$region_mammal) #shows names of brain region levels

levels(data$region_mammal) #shows names of brain region levels

Hipp=subset(data, region_mammal=="Hipp") #makes a dataframe for Hipp

blAMY=subset(data, region_mammal=="blAMY")

LS=subset(data, region_mammal=="LS")

meAMY.BNST=subset(data, region_mammal=="meAMY.BNST")

NAcc=subset(data, region_mammal=="NAcc")

POA=subset(data, region_mammal=="POA")

Str.CP=subset(data, region_mammal=="Str.CP")

VTA=subset(data, region_mammal=="VTA")
```

Do remaining steps for each brain region data frame separately. Here is the example using blAMY

```
# 4. model fitting

table(blAMY$sex, blAMY$social) # shows you table of all combinations of all factos, for N reporting purposes

## specify your "reference" comparisons

blAMY$sex = relevel(blAMY$sex, ref='M')
```

blAMY$social = relevel(blAMY$social, ref='PB')

#### fit naive model (assumes no control genes)

blAMY_naive_model=mcmc.qpcr(data=blAMY, fixed="sex+social+sex:social",random = c("individual"), nitt=510000,thin=500,burnin=10000)

summary(blAMY_naive_model) # this shows results from fully-crossed 2-way fixed effects

HPDsumm_blAMY_naive=HPDsummary(blAMY_naive_model, blAMY, relative=T)  #plot with HDPsummary to visualize whether there are global effects (i.e., all genes appear to be up- or down-regulated under some conditions

### 5. decision time

#### if no global effects are present in naive model, try sharpening credible intervals by running an "informed" model instead, but only if your control genes are reasonably stable in the naive model (as per HPDsymmary plot).

#### if global effects ARE present in naive model, then fit a "soft normalization" model (pg. 30 Matz tutorial, step 10)

##### testing whether informed model sharpens credible intervals more than naive model:

blAMY_naive_model = mcmc.qpcr(data=blAMY, fixed="sex+social+sex:social", random = "individual",nitt=510000,thin=500,burnin=10000, controls=c("r18S"), include=0)

blAMY_informed_model = mcmc.qpcr(data=blAMY, fixed="sex+social+sex:social", random = "individual", controls=c("r18S"), m.fix=1.2, nitt=510000,thin=500,burnin=10000)

HPDplot(model=blAMY_naive_model, factors="socialSol", main="social_system", hpdtype="l")

HPDpoints(model=blAMY_informed_model, factors="socialSol", hpdtype="l", col="coral") #plots credible interval difference between naive and informed

### 6. In the best fit model, conduct pair-wise comparisons of p-values:

blAMY_naive_pwp = HPDsummary(model=blAMY_naive_model, data=blAMY) # shows absolute abundances

blAMY_naive_pwp$summary # retrieves bundles of data that can be used for more plotting (means, sds, CIs)

blAMY_naive_pwp$geneWise # calculates pairwise comparisons

### 7. plot results

plot_blAMY_naive=HPDsummary(blAMY_naive_model, blAMY,xgroup="social") # trellis plot of all genes together in one panel trellisByGene(plot_blAMY_naive, xFactor="social", groupFactor = "sex")+xlab("social system") # trellis plot of genes shown in separate panels

#end

*Chaetodon species: Social system X sex*

### 1. install.packages("MCMC.qpcr")

library(MCMC.qpcr)

### 2. read in data

data <- read.csv(file="Nowicki_C_SPP_M&F_data_ESM.csv")

data$count <- as.numeric(data$count) # force counts to be numeric

head(data) #allows you to view your data to ensure it was imported correctly

### 3. sub-set your data into different brain regions and store them in separate data frames, in order to analyze each brain region separately

levels(data$region_mammal) #shows names of brain region levels

Hipp=subset(data, region_mammal=="Hipp") #makes a dataframe for Hipp

blAMY=subset(data, region_mammal=="blAMY")

LS=subset(data, region_mammal=="LS")

meAMY.BNST=subset(data, region_mammal=="meAMY.BNST")

NAcc=subset(data, region_mammal=="NAcc")

POA=subset(data, region_mammal=="POA")

Str.CP=subset(data, region_mammal=="Str.CP")

VTA=subset(data, region_mammal=="VTA")

Do remaining steps for each brain region data frame separately. Here is the example using blAMY:

### 4. model fitting

table(blAMY$sex, blAMY$social) # shows you table of all combinations of all factos, for N reporting purposes

#### specify your "reference" comparisons

blAMY$sex = relevel(blAMY$sex, ref='M') #sets M as reference for sex factor

blAMY$social = relevel(blAMY$social, ref='PB') #sets PB as reference for social factor

#### fit naive model (assumes no control genes)

blAMY_naive_model=mcmc.qpcr(data=blAMY, fixed="sex+social+sex:social",random = c("individual"), nitt=510000,thin=500,burnin=10000)

summary(blAMY_naive_model) # this shows results from fully-crossed 2-way fixed effects

blAMY_HPDsumm_naive_model=HPDsummary(blAMY_naive_model, blAMY, relative=T) #plot with HDPsummary to visualize whether there are global effects (i.e., all genes appear to be up- or down-regulated under some conditions)

### 5. decision time

#### if no global effects are present in naive model, try sharpening credible intervals by running an "informed" model instead, but only if your control genes are reasonably stable in the naive model (as per HPDsymmary plot).

#### if global effects ARE present in naive model, then fit a "soft normalization" model (pg. 30 tutorial, step 10).

##### testing whether informed model sharpens credible intervals more than naive model

blAMY_naive_model = mcmc.qpcr(data=blAMY, fixed="sex+social+sex:social", random = "individual",nitt=510000,thin=500,burnin=10000, controls=c("r18S"), include=0) #analyzes sex-specific effects of social system

blAMY_informed_model = mcmc.qpcr(data=blAMY, fixed="sex+social+sex:social", random = "individual", controls=c("r18S"), m.fix=1.2, nitt=510000,thin=500,burnin=10000) #analyzes sex-specific effects of social system

HPDplot(model=blAMY_naive_model, factors="socialSol", main="social_system", hpdtype="l") #plots credible interval of naive model

HPDpoints(model=blAMY_informed_model, factors="socialSol", hpdtype="l", col="coral") #plots credible interval difference between naive and informed

### 6. in best fit model, conduct pair-wise comparisons of p-values

blAMY_informed_pwp = HPDsummary(model=blAMY_informed_model, data=blAMY) # shows absolute abundances blAMY_informed_pwp$summary # retrieves bundles of data that can be used for more plotting (means, sds, CIs)

blAMY_informed_pwp$geneWise # calculates pairwise comparisons

### 7. plot results

plot_blAMY_infor=HPDsummary(blAMY_informed_model, blAMY,xgroup="social") # trellis plot of all genes together in same panel

trellisByGene(plot_blAMY_infor, xFactor="social", groupFactor = "sex")+xlab("social system") # trellis plot of each gene in its own panel

#end.

*Chaetodon species: Males: OTR (ITR) in meAMY.BNST (Vs):*

### 1. install.packages("MCMC.qpcr")

library(MCMC.qpcr)

### 2. read in data

data <- read.csv(file="Nowicki_C_SPP_Males_data_ESM.csv")

data$count <- as.numeric(data$count) # force counts to be numeric

head(data) # allows you to view your data to ensure it was imported correctly

levels(data$sex) # shows you all levels of the sex factor, to sanity check the data set looks correct

### 3. sub-set data into different brain regions and store them in separate data frames, in order to analyze each brain region separately

levels(data$region_mammal) #shows names of brain region levels

Hipp=subset(data, region_mammal=="Hipp") #makes a dataframe for Hipp

blAMY=subset(data, region_mammal=="blAMY")

LS=subset(data, region_mammal=="LS")

meAMY.BNST=subset(data, region_mammal=="meAMY.BNST")

NAcc=subset(data, region_mammal=="NAcc")

POA=subset(data, region_mammal=="POA")

```
Str.CP=subset(data, region_mammal=="Str.CP")

VTA=subset(data, region_mammal=="VTA")

# 4. model fitting

meAMY.BNST$species = relevel(meAMY.BNST $species, ref='C.bar') #sets C.bar as "reference” for species factor

meAMY.BNST_naive_model=mcmc.qpcr(data=meAMY.BNST, fixed="species", random = "individual", nitt=510000,thin=500,burnin=10000, pr=TRUE)

summary(meAMY.BNST_naive_model) # this shows results from  1-way fixed effects

meAMY.BNST_HPDsumm_naive_model=HPDsummary(meAMY.BNST_naive_model, meAMY.BNST, relative=T) #eyeballametrically determine whether there are global effects.

# 5. decision time

## if no global effects are present in naive model, try sharpening credible inervals by running an "informed" model instead.

## if global effects ARE present in naive model, then fit a "soft normalization" model (pg. 30 Matz tutorial, step 10)

### testing whether informed model sharpens credible intervals more than naive model:

MeAMY.BNST_naive_model = mcmc.qpcr(data=MeAMY.BNST, fixed="species", random = "individual", controls=c("r18S"), include=0, nitt=510000,thin=500,burnin=10000) #runs naive model

MeAMY.BNST_informed_model = mcmc.qpcr(data=MeAMY.BNST, fixed="species", random = "individual", controls=c("r18S"), m.fix=1.2, nitt=510000,thin=500,burnin=10000) #runs informed model

HPDplot(model=MeAMY.BNST_naive_model, factors="speciesC.lun", main="C.lun", hpdtype="l") #plots credible interval of naive model

HPDpoints(model=MeAMY.BNST_informed_model, factors="speciesC.lun", hpdtype="l", col="coral") #plots credible interval difference between naive and informed

# 6. In the best fit model, conduct pair-wise comparisons of p-values

MeAMY.BNST_spp_informed_pwp = HPDsummary(model=MeAMY.BNST_informed_model, data=MeAMY.BNST) # shows absolute abundances

MeAMY.BNST_spp_informed_pwp$summary # retrieves bundles of data that can be used for more plotting (means, sds, CIs)

MeAMY.BNST_spp_informed_pwp$geneWise # calculates pairwise difference between treatments and their statistical significances; upper triangle is log fold changes, lower triangle is the corresponding p-values
```

```
# 7. plot results

spp_order<-c("C.vag", "C.lun", "C.bar", "C.trif", "C.rainf", "C.pleb")

plot_meAMY.BNST_informed=HPDsummary(meAMY.BNST_informed_model,
meAMY.BNST,xgroup="species", x.order=spp_order) # trellis plot of all genes in same panel

trellisByGene(plot_meAMY.BNST_informed, xFactor="species", groupFactor =
"species")+xlab("species") # trellis plots of genes in separate panels

#end.
```

**Supplementary Table S1**: Primer sequences used to reverse transcribe, clone, and preform quantitative qPCR on *Chaetodon* butterflyfish

| **Target gene** | **Cloning primers** | **Reverse Transcription primers** | **qPCR primers** |
|---|---|---|---|
| ITR | Pair a (using *C.lunulatus* transcriptome):<br>F: 5'-TTTTGTGCAGGTTGGTGAAA<br>R:5'-AGATCCAGGGGTTACAGCAG<br><br>Pair b (to obtain exon sequence):<br>F: 5'- TTTTGTGCAGGTTGGTGAAA<br>R: 5'- GAATTGAACCGCTGGATGTT | R:5'- GGCTGCTCGTGCTTTTAATG | F:5'- GTCTGTTGGACCCCCTTTTT<br>R:5'- CAGCAGCATGGAGATGATGA |
| V1aR | Pair a (using *C.lunulatus* transcriptome):<br>F: 5'-GGAAGACGATGACTGGTGCT<br>R: 5'-AGCTGTTGAGACTGGCAAGG<br><br>Pair b (to obtain exon sequence):<br>F: 5'- GGAAGACGATGACTGGTGCT<br>R: 5'- TGTTAGACCTCCTGGCTGCT | R:5'-AGGGCTTCGATTGGTCATCT | F:5'-CTGTGTGGGATGAAAACTTCCT<br>R:5'-AGGAGGTGACCGCTGAAGAT |
| D1R | Pair a (using *C.lunulatus* transcriptome):<br>F: 5'-GAACGCAAGATGACCCCTAA<br>R: 5'-CCTGTCAGGCATGTCCTTTT | R:5'-TCAAAGGTGGAGCTGAT | F:5'-TCGAACATGGAGAGTGAGAGC<br>R:5-CCAGCAGCACACAAACACTC |
| D2R | Pair a (using *C.lunulatus* transcriptome):<br>F: 5'-TTGCTGTAAGCTGCCATTTG<br>R: 5'-TTTTGCCTGAAACAGGTCA<br><br>Pair b (using *C.lunulatus* transcriptome):<br>F: 5'-TCTGTGGTGTGGGTGCTGT<br>R: 5'-CACGCCTTGCAATAAGACAC | R:5'-TCTGTTGCAGGATCTCCATTC | F:5'-AACGGGAGCTTTCCTGTCA<br>R:5'-GCTGTTGTTCAGCTCATCCAG |
| MOR | Pair a (using *C.lunulatus* transcriptome):<br>F: 5'-AGACCGCCACCAACATCTAC<br>R: 5'-GGATGAGGGTTACGACAGGA<br><br>Pair b (using *C.lunulatus* transcriptome):<br>F: 5'-AGCACGCTACCTTTCCAGAG<br>R: 5'-CCCTTGAAGTTCTCATCCAG | R:5'- CGCAGGTTCCTGTCCTTCT | F:5'- TCATGTTCATGGCCTCCAC<br>R:5'- GCAGATCTTCAGCAGGGTGT |
| 18S | Pair a (using *C.lunulatus* transcriptome):<br>F: 5'-GAGACTCCGGCATGCTAACT<br>R: 5'-GTAATGATCCTTCCGCAGGT | 5'- ATAGTCAAGTTTGATCGTCTTCTCG | F:5'- CAGTAAGCGCGGGTCATAAG<br>R:5'- CGATCCGAGGACCTCACTAA |

**Supplementary Table S2**: Thermal cycle parameters used to preform quantitative qPCR on *Chaetodon* butterflyfish

| Gene | Enzyme mix | Cycle function | No. cycles | Temp. (°C) | Time |
|---|---|---|---|---|---|
| 18S, V1aR, ITR | PerfeCTa SYBR® Green SuperMix | Initial denature | 1 | 95 | 30 sec |
| | | Denature<br>Primer anneal<br>Extension<br>+ Plate read | 45 | 95<br>55<br>70 | 5 sec<br>15 sec<br>10 sec |
| | | Melt curve<br>+ Plate read | 1 | 65-95, 0.5 increment | 5 sec |
| MOR, D1R | PerfeCTa SYBR® Green SuperMix | Initial denature | 1 | 95 | 30 sec |
| | | Denature<br>Primer anneal<br>Extension<br>+ Plate read | 45 | 95<br>60<br>70 | 5 sec<br>15 sec<br>10 sec |
| | | Melt curve<br>+ Plate read | 1 | 65-95, 0.5 increment | 5 sec |
| D2R | PerfeCTa SYBR® Green FastMix | Initial denature | 1 | 95 | 3 min |
| | | Denature<br>Primer anneal + extension<br>+ Plate read | 45 | 95<br>60 | 15 sec<br>1 min |
| | | Melt curve<br>+ Plate read | 1 | 65-95, 0.5 increment | 5 sec |

**Supplementary Table S3:** Model summary of main and interactive effects of sex and social system on brain region-specific gene expression differences in *Chaetodon lunulatus* butterflyfishes. Results are reported as natural log-fold changes from the *a priori* comparison state of male pair bonding, with a two-tailed p value. Associated pair-wise differences between treatments (upper triangle) and their statistical significance (lower triangle) are reported for each brain region and gene, below summary statistics.

| **Dm (blAMY)** | post.mean | l-95% | u-95% | eff.samp | pMCMC |
|---|---|---|---|---|---|
| geneD1R | 3.142 | 1.8472 | 4.5182 | 1000 | <0.001 |
| geneD2R | 3.6135 | 1.732 | 5.4551 | 1000 | <0.001 |
| geneITR | -2.7655 | -5.7253 | 0.1035 | 194.148 | 0.016 |
| geneMOR | 2.5017 | 0.9068 | 4.0094 | 895.047 | 0.004 |
| gener18S | 14.9415 | 13.5903 | 16.2004 | 1000 | <0.001 |
| geneV1aR | -2.3168 | -4.7494 | -0.129 | 858.875 | 0.03 |
| geneD1R:sexF | 1.3124 | -0.8307 | 3.4848 | 1000 | 0.248 |
| geneD2R:sexF | -1.4817 | -3.9562 | 1.4242 | 754.521 | 0.252 |
| geneITR:sexF | -1.9483 | -6.3603 | 2.6047 | 693.048 | 0.366 |
| geneMOR:sexF | 0.3657 | -1.9418 | 2.7032 | 1000 | 0.758 |
| gener18S:sexF | -0.6788 | -2.5982 | 1.1364 | 1000 | 0.46 |
| geneV1aR:sexF | 1.1761 | -1.8762 | 3.8531 | 1000 | 0.396 |
| geneD1R:socialSol | -1.4655 | -3.5313 | 0.8352 | 1000 | 0.176 |
| geneD2R:socialSol | -1.2047 | -3.6959 | 1.8502 | 1096.457 | 0.382 |
| geneITR:socialSol | -193.962 | -357.689 | -5.2094 | 1.287 | 0.004 |
| geneMOR:socialSol | -2.9367 | -5.4574 | -0.5429 | 892.836 | 0.014 |
| gener18S:socialSol | -0.6569 | -2.7192 | 1.4031 | 777.435 | 0.498 |
| geneV1aR:socialSol | -1.4491 | -5.0021 | 2.1111 | 1000 | 0.422 |
| geneD1R:sexF:socialSol | -1.2232 | -4.1923 | 2.2079 | 1000 | 0.428 |
| geneD2R:sexF:socialSol | 1.8539 | -1.6811 | 6.3018 | 1000 | 0.344 |
| geneITR:sexF:socialSol | 54.4327 | -139.685 | 217.4457 | 4.289 | 0.454 |
| geneMOR:sexF:socialSol | 1.3268 | -2.0225 | 4.6156 | 1000 | 0.436 |
| gener18S:sexF:socialSol | 0.7555 | -2.1329 | 3.6111 | 1000 | 0.628 |
| geneV1aR:sexF:socialSol | -1.1577 | -6.1427 | 4.6096 | 891.465 | 0.644 |

**Dm (blAMY) Pair-wise p values**

| | | | | | |
|---|---|---|---|---|---|
| $D1R | | | | | |
| | difference | | | | |
| pvalue | | sexM:socialPB | sexM:socialSol | sexF:socialPB | sexF:socialSol |
| | sexM:socialPB | NA | -2.11433467 | 1.89332498 | -1.985673 |
| | sexM:socialSol | 0.191114 | NA | 4.00765965 | 0.1286617 |
| | sexF:socialPB | 0.2548586 | 0.02636481 | NA | -3.878998 |
| | sexF:socialSol | 0.1915565 | 0.93752835 | 0.02801988 | NA |
| $D2R | | | | | |
| | difference | | | | |
| pvalue | | sexM:socialPB | sexM:socialSol | sexF:socialPB | sexF:socialSol |
| | sexM:socialPB | NA | -1.7380322 | -2.1376927 | -1.2011756 |
| | sexM:socialSol | 0.3943747 | NA | -0.3996604 | 0.5368567 |
| | sexF:socialPB | 0.2741771 | 0.8453605 | NA | 0.9365171 |
| | sexF:socialSol | 0.5502811 | 0.8062066 | 0.6562063 | NA |
| $ITR | | | | | |
| | difference | | | | |

| | | | | | |
|---|---|---|---|---|---|
| pvalue | | sexM:socialPB | sexM:socialSol | sexF:socialPB | sexF:socialSol |
| | sexM:socialPB | NA | -279.8274998 | -2.8108276 | -204.10856 |
| | sexM:socialSol | 0.1066488 | NA | 277.0166723 | 75.71894 |
| | sexF:socialPB | 0.43701 | 0.110682 | NA | -201.29773 |
| | sexF:socialSol | 0.1361658 | 0.5404262 | 0.1417471 | NA |
| $MOR | | | | | |
| | difference | | | | |
| pvalue | | sexM:socialPB | sexM:socialSol | sexF:socialPB | sexF:socialSol |
| | sexM:socialPB | NA | -4.236693617 | 0.5275561 | -1.794922 |
| | sexM:socialSol | 0.01897804 | NA | 4.7642498 | 2.441772 |
| | sexF:socialPB | 0.75241678 | 0.009653445 | NA | -2.322478 |
| | sexF:socialSol | 0.30748278 | 0.188131608 | 0.1757757 | NA |
| $r18S | | | | | |
| | difference | | | | |
| pvalue | | sexM:socialPB | sexM:socialSol | sexF:socialPB | sexF:socialSol |
| | sexM:socialPB | NA | -0.9477639 | -0.97925093 | -0.8370861 |
| | sexM:socialSol | 0.5288687 | NA | -0.03148701 | 0.1106778 |
| | sexF:socialPB | 0.4840288 | 0.983036 | NA | 0.1421648 |
| | sexF:socialSol | 0.5796442 | 0.9463554 | 0.92497113 | NA |
| $V1aR | | | | | |
| | difference | | | | |
| pvalue | | sexM:socialPB | sexM:socialSol | sexF:socialPB | sexF:socialSol |
| | sexM:socialPB | NA | -2.0906618 | 1.6968106 | -2.064007 |
| | sexM:socialSol | 0.4571102 | NA | 3.7874725 | 0.02665483 |
| | sexF:socialPB | 0.4261231 | 0.1733417 | NA | -3.76081764 |
| | sexF:socialSol | 0.4656879 | 0.9937238 | 0.1794475 | NA |

| **DI (Hipp)** | post.mean | l-95% | u-95% | eff.samp | pMCMC |
|---|---|---|---|---|---|
| geneD1R | 4.32E+00 | 2.29E+00 | 6.54E+00 | 1215.32 | <0.001 |
| geneD2R | 2.16E+00 | -2.01E-01 | 4.38E+00 | 752.78 | 0.086 |
| geneITR | -3.25E+00 | -6.07E+00 | -7.73E-01 | 1000 | 0.004 |
| geneMOR | 2.98E+00 | 1.24E+00 | 4.73E+00 | 1000 | 0.002 |
| gener18S | 1.40E+01 | 1.23E+01 | 1.55E+01 | 1000 | <0.001 |
| geneV1aR | -2.49E+00 | -4.79E+00 | -4.25E-02 | 875.43 | 0.024 |
| geneD1R:sexF | -8.71E-01 | -3.31E+00 | 2.10E+00 | 1122.67 | 0.508 |
| geneD2R:sexF | 1.54E+00 | -1.73E+00 | 4.64E+00 | 809.55 | 0.304 |
| geneITR:sexF | 2.46E+00 | -3.04E-01 | 5.86E+00 | 1000 | 0.11 |
| geneMOR:sexF | -8.96E-01 | -3.29E+00 | 1.99E+00 | 897.13 | 0.468 |
| gener18S:sexF | 1.18E+00 | -9.17E-01 | 3.53E+00 | 1000 | 0.298 |
| geneV1aR:sexF | 1.36E+00 | -1.88E+00 | 4.47E+00 | 1000 | 0.358 |
| geneD1R:socialSol | -2.20E+00 | -5.29E+00 | 4.48E-01 | 1000 | 0.128 |
| geneD2R:socialSol | -6.48E-03 | -3.33E+00 | 3.16E+00 | 1134.11 | 0.958 |
| geneITR:socialSol | -1.09E+00 | -5.24E+00 | 3.52E+00 | 1000 | 0.634 |
| geneMOR:socialSol | -1.36E+00 | -4.07E+00 | 1.48E+00 | 1164.88 | 0.318 |
| gener18S:socialSol | 2.29E-01 | -2.15E+00 | 2.66E+00 | 1000 | 0.846 |
| geneV1aR:socialSol | -2.19E+00 | -6.65E+00 | 2.00E+00 | 1000 | 0.286 |
| geneD1R:sexF:socialSol | 4.05E-01 | -3.41E+00 | 4.13E+00 | 1000 | 0.8 |
| geneD2R:sexF:socialSol | -1.06E+00 | -5.49E+00 | 3.38E+00 | 1000 | 0.642 |

| | | | | | |
|---|---|---|---|---|---|
| geneITR:sexF:socialSol | 7.91E-01 | -4.52E+00 | 5.26E+00 | 1000 | 0.734 |
| geneMOR:sexF:socialSol | 1.50E+00 | -2.30E+00 | 5.47E+00 | 1000 | 0.45 |
| gener18S:sexF:socialSol | -1.31E+00 | -4.79E+00 | 1.87E+00 | 890.17 | 0.41 |
| geneV1aR:sexF:socialSol | -6.29E+01 | -1.55E+02 | -4.37E-01 | 5.88 | 0.012 |

**DI (Hipp) Pair-wise p values**

| | | | | | |
|---|---|---|---|---|---|
| $D1R | | | | | |
| | difference | | | | |
| pvalue | | sexM:socialPB | sexM:socialSol | sexF:socialPB | sexF:socialSol |
| | sexM:socialPB | NA | -3.16841 | -1.25707 | -3.840596 |
| | sexM:socialSol | 0.134117 | NA | 1.911336 | -0.6721889 |
| | sexF:socialPB | 0.520008 | 0.295636 | NA | -2.5835252 |
| | sexF:socialSol | 0.057376 | 0.715174 | 0.130778 | NA |
| $D2R | | | | | |
| | difference | | | | |
| pvalue | | sexM:socialPB | sexM:socialSol | sexF:socialPB | sexF:socialSol |
| | sexM:socialPB | NA | -0.00935 | 2.224183 | 0.6843199 |
| | sexM:socialSol | 0.996827 | NA | 2.233538 | 0.6936748 |
| | sexF:socialPB | 0.339648 | 0.352695 | NA | -1.539863 |
| | sexF:socialSol | 0.765696 | 0.780998 | 0.513554 | NA |
| $ITR | | | | | |
| | difference | | | | |
| pvalue | sexM:socialPB | sexM:socialSol | sexF:socialPB | sexF:socialSol | |
| | sexM:socialPB | NA | -1.57588 | 3.553385 | 3.1190451 |
| | sexM:socialSol | 0.62438 | NA | 5.129269 | 4.6949293 |
| | sexF:socialPB | 0.124599 | 0.0759 | NA | -0.4343394 |
| | sexF:socialSol | 0.198681 | 0.107248 | 0.828115 | NA |
| $MOR | | | | | |
| | difference | | | | |
| pvalue | | sexM:socialPB | sexM:socialSol | sexF:socialPB | sexF:socialSol |
| | sexM:socialPB | NA | -1.96844 | -1.2933 | -1.0965406 |
| | sexM:socialSol | 0.335637 | NA | 0.675135 | 0.8718954 |
| | sexF:socialPB | 0.503268 | 0.746256 | NA | 0.1967603 |
| | sexF:socialSol | 0.576531 | 0.687927 | 0.920763 | NA |
| $r18S | | | | | |
| | difference | | | | |
| pvalue | | sexM:socialPB | sexM:socialSol | sexF:socialPB | sexF:socialSol |
| | sexM:socialPB | NA | 0.330311 | 1.701266 | 0.1366222 |
| | sexM:socialSol | 0.85299 | NA | 1.370955 | -0.1936886 |
| | sexF:socialPB | 0.303461 | 0.435768 | NA | -1.5646433 |
| | sexF:socialSol | 0.933051 | 0.912509 | 0.350976 | NA |
| $V1aR | | | | | |

| | difference | | | | |
|---|---|---|---|---|---|
| pvalue | | sexM:socialPB | sexM:socialSol | sexF:socialPB | sexF:socialSol |
| | sexM:socialPB | NA | -3.15191 | 1.954299 | -91.86627 |
| | sexM:socialSol | 0.318081 | NA | 5.106213 | -88.71436 |
| | sexF:socialPB | 0.388011 | 0.101299 | NA | -93.82057 |
| | sexF:socialSol | 0.147846 | 0.162791 | 0.138922 | NA |

| **Vv/Vl (LS)** | post.mean | l-95% | u-95% | eff.samp | pMCMC |
|---|---|---|---|---|---|
| geneD1R | 4.18263 | 2.05947 | 6.37721 | 1000 | <0.001 |
| geneD2R | 3.43398 | 1.6286 | 5.15397 | 1000 | <0.001 |
| geneITR | -1.58519 | -4.15107 | 0.43111 | 1000 | 0.158 |
| geneMOR | 2.5583 | 0.52161 | 4.40675 | 1000 | 0.012 |
| geneV1aR | 0.80748 | -1.27265 | 2.80046 | 1000 | 0.434 |
| gener18S | 12.12274 | 10.80329 | 13.49123 | 1000 | <0.001 |
| geneD1R:sexF | -0.10344 | -3.66066 | 3.3877 | 1000 | 0.98 |
| geneD2R:sexF | 0.61094 | -2.46058 | 3.68002 | 1000 | 0.692 |
| geneITR:sexF | -0.55273 | -4.16377 | 3.09086 | 1000 | 0.75 |
| geneMOR:sexF | 0.1833 | -3.10986 | 3.36018 | 1000 | 0.924 |
| geneV1aR:sexF | -1.77035 | -5.84196 | 1.59303 | 990.1 | 0.346 |
| gener18S:sexF | -0.02783 | -0.95233 | 1.11198 | 1000 | 0.962 |
| geneD1R:socialSol | -1.80785 | -4.79607 | 1.31936 | 1000 | 0.242 |
| geneD2R:socialSol | 0.3149 | -2.46907 | 3.3038 | 1000 | 0.806 |
| geneITR:socialSol | 0.49331 | -2.49085 | 3.68865 | 864.6 | 0.734 |
| geneMOR:socialSol | 0.52584 | -2.27097 | 3.28731 | 1000 | 0.752 |
| geneV1aR:socialSol | -3.04181 | -6.09046 | 0.1176 | 1000 | 0.052 |
| gener18S:socialSol | 0.44774 | -0.66276 | 1.48162 | 1127.8 | 0.406 |
| geneD1R:sexF:socialSol | 0.12635 | -4.26795 | 4.60353 | 1000 | 0.964 |
| geneD2R:sexF:socialSol | -1.2808 | -5.1135 | 3.1451 | 1000 | 0.508 |
| geneITR:sexF:socialSol | 0.5128 | -4.53174 | 5.0883 | 907.2 | 0.832 |
| geneMOR:sexF:socialSol | -0.15703 | -4.19607 | 4.37666 | 1000 | 0.934 |
| geneV1aR:sexF:socialSol | 3.1815 | -1.61693 | 8.2854 | 961.6 | 0.192 |
| gener18S:sexF:socialSol | 0.33206 | -0.73743 | 1.47798 | 1095.3 | 0.552 |

**Vv/vl (LS) Pair-wise p values**

| $D1R | | | | | |
|---|---|---|---|---|---|
| | difference | | | | |
| pvalue | | sexM:socialPB | sexM:socialSol | sexF:socialPB | sexF:socialSol |
| | sexM:socialPB | NA | -2.60817 | -0.14923 | -2.575114 |
| | sexM:socialSol | 0.23875 | NA | 2.458942 | 0.03305623 |
| | sexF:socialPB | 0.954749 | 0.361308 | NA | -2.4258854 |
| | sexF:socialSol | 0.232368 | 0.987686 | 0.342613 | NA |
| $D2R | | | | | |
| | difference | | | | |
| pvalue | | sexM:socialPB | sexM:socialSol | sexF:socialPB | sexF:socialSol |
| | sexM:socialPB | NA | 0.454304 | 0.881397 | -0.5121025 |

| | | | | | |
|---|---|---|---|---|---|
| | sexM:socialSol | 0.824989 | NA | 0.427094 | -0.9664061 |
| | sexF:socialPB | 0.701724 | 0.862737 | NA | -1.3934996 |
| | sexF:socialSol | 0.793223 | 0.641384 | 0.535239 | NA |
| $ITR | | | | | |
| | difference | | | | |
| pvalue | | sexM:socialPB | sexM:socialSol | sexF:socialPB | sexF:socialSol |
| | sexM:socialPB | NA | 0.711696 | -0.79742 | 0.65409772 |
| | sexM:socialSol | 0.763474 | NA | -1.50911 | -0.0575984 |
| | sexF:socialPB | 0.770038 | 0.584709 | NA | 1.45151566 |
| | sexF:socialSol | 0.778172 | 0.980471 | 0.586629 | NA |
| $MOR | | | | | |
| | difference | | | | |
| pvalue | | sexM:socialPB | sexM:socialSol | sexF:socialPB | sexF:socialSol |
| | sexM:socialPB | NA | 0.758632 | 0.264452 | 0.79653296 |
| | sexM:socialSol | 0.724684 | NA | -0.49418 | 0.03790077 |
| | sexF:socialPB | 0.912191 | 0.834256 | NA | 0.53208102 |
| | sexF:socialSol | 0.695443 | 0.985817 | 0.819464 | NA |
| $V1aR | | | | | |
| | difference | | | | |
| pvalue | | sexM:socialPB | sexM:socialSol | sexF:socialPB | sexF:socialSol |
| | sexM:socialPB | NA | -4.38841 | -2.55408 | -2.3525538 |
| | sexM:socialSol | 0.058951 | NA | 1.834325 | 2.0358532 |
| | sexF:socialPB | 0.353328 | 0.541294 | NA | 0.2015281 |
| | sexF:socialSol | 0.270102 | 0.426806 | 0.942783 | NA |
| $r18S | | | | | |
| | difference | | | | |
| pvalue | | sexM:socialPB | sexM:socialSol | sexF:socialPB | sexF:socialSol |
| | sexM:socialPB | NA | 0.645945 | -0.04014 | 1.0848603 |
| | sexM:socialSol | 0.409982 | NA | -0.68609 | 0.4389151 |
| | sexF:socialPB | 0.957733 | 0.540707 | NA | 1.1250035 |
| | sexF:socialSol | 0.371343 | 0.675555 | 0.284979 | NA |

| **Vs (meAMY/BNST)** | post.mean | l-95% | u-95% | eff.samp | pMCMC |
|---|---|---|---|---|---|
| geneD1R | 0.41519 | -1.73372 | 2.22517 | 1000 | 0.62 |
| geneD2R | 2.66415 | 0.95883 | 4.51436 | 1000 | 0.008 |
| geneITR | -2.7834 | -5.40423 | 0.06744 | 286 | 0.028 |
| geneMOR | 2.0634 | -2.00066 | 5.88315 | 1000 | 0.28 |
| gener18S | 13.8691 | 12.08308 | 15.4984 | 1000 | <0.001 |
| geneV1aR | -0.66775 | -2.56781 | 1.55733 | 1351.4 | 0.514 |
| geneD1R:sexF | 4.87635 | 1.9087 | 7.92413 | 1000 | 0.006 |
| geneD2R:sexF | 0.29534 | -2.23421 | 3.10785 | 1000 | 0.82 |
| geneITR:sexF | 1.3332 | -1.86136 | 4.90466 | 766.5 | 0.442 |
| geneMOR:sexF | 0.48468 | -4.99799 | 5.59678 | 1052 | 0.818 |

| | | | | | |
|---|---|---|---|---|---|
| gener18S:sexF | -0.33156 | -2.85742 | 2.0727 | 1000 | 0.796 |
| geneV1aR:sexF | 0.32751 | -2.61109 | 3.25598 | 1000 | 0.836 |
| geneD1R:socialSol | 2.12168 | -0.79227 | 5.04754 | 1000 | 0.16 |
| geneD2R:socialSol | 0.08216 | -2.70677 | 3.24178 | 753.5 | 0.964 |
| geneITR:socialSol | 0.43815 | -3.59942 | 4.77041 | 804.2 | 0.856 |
| geneMOR:socialSol | 1.33059 | -4.56275 | 7.17892 | 1000 | 0.654 |
| gener18S:socialSol | -0.41375 | -3.33612 | 2.48778 | 855 | 0.762 |
| geneV1aR:socialSol | -0.79005 | -3.89873 | 2.71125 | 1000 | 0.644 |
| geneD1R:sexF:socialSol | -4.64349 | -8.95856 | -0.44833 | 1000 | 0.032 |
| geneD2R:sexF:socialSol | -0.54687 | -4.28436 | 3.76456 | 1000 | 0.752 |
| geneITR:sexF:socialSol | -1.14534 | -6.60084 | 3.9961 | 680.6 | 0.668 |
| geneMOR:sexF:socialSol | -0.43071 | -8.13261 | 6.69024 | 1000 | 0.89 |
| gener18S:sexF:socialSol | 1.13119 | -2.8092 | 5.03306 | 1000 | 0.55 |
| geneV1aR:sexF:socialSol | -1.32989 | -6.23568 | 3.10742 | 1000 | 0.568 |

**Vv/vl (LS) Pair-wise p values**

| | | | | | |
|---|---|---|---|---|---|
| $D1R | | | | | |
| | difference | | | | |
| pvalue | | sexM:socialPB | sexM:socialSol | sexF:socialPB | sexF:socialSol |
| | sexM:socialPB | NA | 3.060942 | 7.035079 | 3.3968875 |
| | sexM:socialSol | 0.162169 | NA | 3.974138 | 0.3359458 |
| | sexF:socialPB | 0.001695 | 0.099793 | NA | -3.6381918 |
| | sexF:socialSol | 0.080558 | 0.876204 | 0.094788 | NA |
| $D2R | | | | | |
| | difference | | | | |
| pvalue | | sexM:socialPB | sexM:socialSol | sexF:socialPB | sexF:socialSol |
| | sexM:socialPB | NA | 0.118536 | 0.42608 | -0.2443452 |
| | sexM:socialSol | 0.955139 | NA | 0.307544 | -0.3628815 |
| | sexF:socialPB | 0.825598 | 0.885753 | NA | -0.670425 |
| | sexF:socialSol | 0.897107 | 0.869571 | 0.728497 | NA |
| $ITR | | | | | |
| | difference | | | | |
| pvalue | | sexM:socialPB | sexM:socialSol | sexF:socialPB | sexF:socialSol |
| | sexM:socialPB | NA | 0.63212 | 1.923402 | 0.9031425 |
| | sexM:socialSol | 0.844609 | NA | 1.291282 | 0.2710224 |
| | sexF:socialPB | 0.489653 | 0.675387 | NA | -1.0202591 |
| | sexF:socialSol | 0.75002 | 0.925349 | 0.71304 | NA |
| $MOR | | | | | |
| | difference | | | | |
| pvalue | | sexM:socialPB | sexM:socialSol | sexF:socialPB | sexF:socialSol |
| | sexM:socialPB | NA | 1.919634 | 0.699246 | 1.99749782 |
| | sexM:socialSol | 0.65115 | NA | -1.22039 | 0.07786436 |
| | sexF:socialPB | 0.855757 | 0.765798 | NA | 1.29825188 |

| | | | | | |
|---|---|---|---|---|---|
| | sexF:socialSol | 0.608205 | 0.983895 | 0.723613 | NA |
| $r18S | | | | | |
| | difference | | | | |
| pvalue | | sexM:socialPB | sexM:socialSol | sexF:socialPB | sexF:socialSol |
| | sexM:socialPB | NA | -0.59691 | -0.47833 | 0.556718 |
| | sexM:socialSol | 0.782062 | NA | 0.118579 | 1.15363 |
| | sexF:socialPB | 0.795532 | 0.956062 | NA | 1.035051 |
| | sexF:socialSol | 0.7601 | 0.584902 | 0.580568 | NA |
| $V1aR | | | | | |
| | difference | | | | |
| pvalue | | sexM:socialPB | sexM:socialSol | sexF:socialPB | sexF:socialSol |
| | sexM:socialPB | NA | -1.13979 | 0.472503 | -2.585912 |
| | sexM:socialSol | 0.65074 | NA | 1.612297 | -1.446117 |
| | sexF:socialPB | 0.825622 | 0.536849 | NA | -3.058415 |
| | sexF:socialSol | 0.282817 | 0.591447 | 0.212108 | NA |

| **Vd (NAcc)** | post.mean | l-95% | u-95% | eff.samp | pMCMC |
|---|---|---|---|---|---|
| geneD1R | 3.40928 | 1.41427 | 5.25825 | 1000 | <0.001 |
| geneD2R | 2.08561 | 0.45417 | 3.77862 | 1097.557 | 0.02 |
| geneITR | -77.3637 | -160.431 | -5.46338 | 4.538 | <0.001 |
| geneMOR | -34.7492 | -100.592 | 5.37739 | 21.038 | 0.078 |
| gener18S | 12.72855 | 11.57857 | 14.02614 | 1000 | <0.001 |
| geneV1aR | -11.2197 | -40.2993 | 0.08154 | 19.051 | 0.01 |
| geneD1R:sexF | -0.25354 | -2.35008 | 2.08043 | 1000 | 0.798 |
| geneD2R:sexF | 0.51131 | -2.20217 | 2.96635 | 1000 | 0.65 |
| geneITR:sexF | 70.11619 | -3.01837 | 154.5589 | 4.295 | 0.018 |
| geneMOR:sexF | 11.61996 | -66.1972 | 85.19842 | 59.527 | 0.73 |
| gener18S:sexF | -0.17443 | -1.90647 | 1.69883 | 1229.386 | 0.832 |
| geneV1aR:sexF | 6.64009 | -8.94948 | 33.84802 | 37.513 | 0.262 |
| geneD1R:socialSol | -3.63525 | -6.44676 | -0.7174 | 1303.857 | 0.016 |
| geneD2R:socialSol | -2.46742 | -6.27528 | 0.85833 | 1000 | 0.152 |
| geneITR:socialSol | -71.8958 | -256.436 | 91.4815 | 3.502 | 0.662 |
| geneMOR:socialSol | 15.65856 | -75.4037 | 95.17179 | 70.272 | 0.61 |
| gener18S:socialSol | -1.62641 | -3.95243 | 0.54302 | 1000 | 0.144 |
| geneV1aR:socialSol | 6.18767 | -10.2903 | 36.39662 | 81.08 | 0.424 |
| geneD1R:sexF:socialSol | -0.49672 | -3.98204 | 3.29604 | 1000 | 0.776 |
| geneD2R:sexF:socialSol | -0.2799 | -4.57304 | 4.71507 | 1192.098 | 0.87 |
| geneITR:sexF:socialSol | -7.671 | -226.221 | 141.0776 | 3.607 | 0.842 |
| geneMOR:sexF:socialSol | -13.276 | -126.788 | 105.7283 | 52.912 | 0.816 |
| gener18S:sexF:socialSol | 0.80758 | -2.08886 | 3.62423 | 910.999 | 0.54 |
| geneV1aR:sexF:socialSol | -11.9915 | -49.4752 | 10.41948 | 32.394 | 0.184 |

**Vd (NAcc) Pair-wise p values**

| | | | | | |
|---|---|---|---|---|---|
| $D1R | | | | | |
| | difference | | | | |
| pvalue | | sexM:socialPB | sexM:socialSol | sexF:socialPB | sexF:socialSol |
| | sexM:socialPB | NA | -5.24456 | -0.36578 | -6.32695 |
| | sexM:socialSol | 0.011822 | NA | 4.878775 | -1.0824 |
| | sexF:socialPB | 0.824113 | 0.007921 | NA | -5.96117 |
| | sexF:socialSol | 0.000638 | 0.597229 | 0.000169 | NA |
| $D2R | | | | | |
| | difference | | | | |
| pvalue | | sexM:socialPB | sexM:socialSol | sexF:socialPB | sexF:socialSol |
| | sexM:socialPB | NA | -3.55973 | 0.737671 | -3.22586 |
| | sexM:socialSol | 0.181647 | NA | 4.297402 | 0.333867 |
| | sexF:socialPB | 0.696574 | 0.110721 | NA | -3.96354 |
| | sexF:socialSol | 0.078372 | 0.900303 | 0.041888 | NA |
| $ITR | | | | | |
| | difference | | | | |
| pvalue | | sexM:socialPB | sexM:socialSol | sexF:socialPB | sexF:socialSol |
| | sexM:socialPB | NA | -103.724 | 101.1563 | -13.6344 |
| | sexM:socialSol | 0.48642 | NA | 204.88 | 90.08936 |
| | sexF:socialPB | 0.140002 | 0.131735 | NA | -114.791 |
| | sexF:socialSol | 0.809516 | 0.486853 | 0.065646 | NA |
| $MOR | | | | | |
| | difference | | | | |
| pvalue | | sexM:socialPB | sexM:socialSol | sexF:socialPB | sexF:socialSol |
| | sexM:socialPB | NA | 22.59053 | 16.76406 | 20.20134 |
| | sexM:socialSol | 0.700638 | NA | -5.82647 | -2.38919 |
| | sexF:socialPB | 0.7558 | 0.925631 | NA | 3.43728 |
| | sexF:socialSol | 0.645777 | 0.965382 | 0.946784 | NA |
| $r18S | | | | | |
| | difference | | | | |
| pvalue | | sexM:socialPB | sexM:socialSol | sexF:socialPB | sexF:socialSol |
| | sexM:socialPB | NA | -2.34641 | -0.25165 | -1.43296 |
| | sexM:socialSol | 0.153677 | NA | 2.094758 | 0.913447 |
| | sexF:socialPB | 0.845353 | 0.216417 | NA | -1.18131 |
| | sexF:socialSol | 0.251057 | 0.576735 | 0.355302 | NA |
| $V1aR | | | | | |
| | difference | | | | |
| pvalue | | sexM:socialPB | sexM:socialSol | sexF:socialPB | sexF:socialSol |
| | sexM:socialPB | NA | 8.926924 | 9.579622 | 1.206478 |
| | sexM:socialSol | 0.602429 | NA | 0.652698 | -7.72045 |
| | sexF:socialPB | 0.566705 | 0.965746 | NA | -8.37314 |
| | sexF:socialSol | 0.932346 | 0.591932 | 0.513438 | NA |

| **POA (POA)** | post.mean | l-95% | u-95% | eff.samp | pMCMC |
|---|---|---|---|---|---|
| geneD1R | 4.3673 | 2.6354 | 6.1356 | 1000 | <0.001 |
| geneD2R | 3.8541 | 2.2052 | 5.5648 | 1000 | <0.001 |
| geneITR | -0.5018 | -2.1461 | 1.2122 | 904.4 | 0.594 |
| geneMOR | 1.9263 | -0.1851 | 3.7134 | 1000 | 0.056 |
| geneV1aR | -1.2953 | -3.3395 | 0.659 | 1031.2 | 0.212 |
| gener18S | 13.06 | 11.929 | 14.1965 | 1000 | <0.001 |
| geneD1R:sexF | 0.8944 | -1.4561 | 3.1279 | 1000 | 0.466 |
| geneD2R:sexF | 1.2108 | -0.9323 | 3.3259 | 1000 | 0.29 |
| geneITR:sexF | 0.8153 | -1.3922 | 3.1205 | 1000 | 0.52 |
| geneMOR:sexF | 2.4464 | -0.4033 | 4.8277 | 1000 | 0.068 |
| geneV1aR:sexF | 2.7253 | 0.1442 | 5.4334 | 1000 | 0.04 |
| gener18S:sexF | 0.3687 | -0.5607 | 1.4214 | 1000 | 0.46 |
| geneD1R:socialSol | -3.2855 | -6.2908 | -0.6255 | 1000 | 0.028 |
| geneD2R:socialSol | -1.339 | -3.7868 | 1.2976 | 1096 | 0.3 |
| geneITR:socialSol | -1.7395 | -4.4291 | 1.3362 | 1000 | 0.222 |
| geneMOR:socialSol | 0.4157 | -2.3244 | 3.0698 | 1000 | 0.75 |
| geneV1aR:socialSol | 0.1232 | -2.463 | 3.7451 | 1000 | 0.938 |
| gener18S:socialSol | -0.3185 | -1.3934 | 0.5331 | 1000 | 0.53 |
| geneD1R:sexF:socialSol | 1.6993 | -2.0011 | 5.0748 | 1195.5 | 0.356 |
| geneD2R:sexF:socialSol | 0.468 | -3.2633 | 3.681 | 1258.1 | 0.782 |
| geneITR:sexF:socialSol | 1.8269 | -1.978 | 5.4142 | 1000 | 0.344 |
| geneMOR:sexF:socialSol | -0.5078 | -3.8877 | 3.2985 | 1000 | 0.786 |
| geneV1aR:sexF:socialSol | -0.9352 | -4.7723 | 2.5485 | 1000 | 0.662 |
| gener18S:sexF:socialSol | 0.2283 | -0.7923 | 1.3007 | 1000 | 0.66 |

**POA (POA) Pair-wise p values**

| | | | | | |
|---|---|---|---|---|---|
| $D1R | | | | | |
| | difference | | | | |
| pvalue | | sexM:socialPB | sexM:socialSol | sexF:socialPB | sexF:socialSol |
| | sexM:socialPB | NA | -4.74004 | 1.290324 | -0.99813 |
| | sexM:socialSol | 0.024582 | NA | 6.0303639 | 3.741906 |
| | sexF:socialPB | 0.448421 | 0.002152 | NA | -2.28846 |
| | sexF:socialSol | 0.581531 | 0.057079 | 0.1702788 | NA |
| $D2R | | | | | |
| | difference | | | | |
| pvalue | | sexM:socialPB | sexM:socialSol | sexF:socialPB | sexF:socialSol |
| | sexM:socialPB | NA | -1.93176 | 1.7468499 | 0.490215 |
| | sexM:socialSol | 0.311711 | NA | 3.6786112 | 2.421976 |
| | sexF:socialPB | 0.278293 | 0.031272 | NA | -1.25664 |
| | sexF:socialSol | 0.775446 | 0.190447 | 0.4355411 | NA |
| $ITR | | | | | |
| | difference | | | | |

| pvalue | | sexM:socialPB | sexM:socialSol | sexF:socialPB | sexF:socialSol |
|---|---|---|---|---|---|
| | sexM:socialPB | NA | -2.50952 | 1.1762082 | 1.302365 |
| | sexM:socialSol | 0.224518 | NA | 3.6857259 | 3.811883 |
| | sexF:socialPB | 0.498062 | 0.073931 | NA | 0.126157 |
| | sexF:socialSol | 0.472984 | 0.079209 | 0.9425484 | NA |
| $MOR | | | | | |
| | difference | | | | |
| pvalue | | sexM:socialPB | sexM:socialSol | sexF:socialPB | sexF:socialSol |
| | sexM:socialPB | NA | 0.599729 | 3.5294268 | 3.396511 |
| | sexM:socialSol | 0.765904 | NA | 2.9296983 | 2.796782 |
| | sexF:socialPB | 0.059922 | 0.100699 | NA | -0.13292 |
| | sexF:socialSol | 0.067231 | 0.136976 | 0.9366706 | NA |
| $V1aR | | | | | |
| | difference | | | | |
| pvalue | | sexM:socialPB | sexM:socialSol | sexF:socialPB | sexF:socialSol |
| | sexM:socialPB | NA | 0.177783 | 3.9317056 | 2.760335 |
| | sexM:socialSol | 0.936343 | NA | 3.7539222 | 2.582551 |
| | sexF:socialPB | 0.04414 | 0.073261 | NA | -1.17137 |
| | sexF:socialSol | 0.180124 | 0.217466 | 0.5176704 | NA |
| $r18S | | | | | |
| | difference | | | | |
| pvalue | | sexM:socialPB | sexM:socialSol | sexF:socialPB | sexF:socialSol |
| | sexM:socialPB | NA | -0.45956 | 0.5318711 | 0.401606 |
| | sexM:socialSol | 0.528481 | NA | 0.9914331 | 0.861168 |
| | sexF:socialPB | 0.463492 | 0.329933 | NA | -0.13027 |
| | sexF:socialSol | 0.727028 | 0.383092 | 0.8933919 | NA |

| **Vc (Str/CP)** | post.mean | l-95% | u-95% | eff.samp | pMCMC |
|---|---|---|---|---|---|
| geneD1R | 3.83523 | 1.73107 | 6.17979 | 1000 | 0.004 |
| geneD2R | 3.08301 | 1.39313 | 4.83215 | 1024.111 | 0.004 |
| geneITR | -0.66475 | -2.66405 | 1.2628 | 1000 | 0.502 |
| geneMOR | 0.02413 | -4.56867 | 3.78887 | 707.354 | 0.924 |
| gener18S | 12.72933 | 10.73039 | 14.39361 | 901.265 | <0.001 |
| geneV1aR | -2.39528 | -7.21094 | 1.97533 | 487.252 | 0.192 |
| geneD1R:sexF | -0.28664 | -2.97676 | 2.8047 | 1000 | 0.83 |
| geneD2R:sexF | 1.32038 | -1.04093 | 3.61791 | 1073.205 | 0.294 |
| geneITR:sexF | -0.66109 | -3.3408 | 2.2858 | 1000 | 0.694 |
| geneMOR:sexF | 0.8442 | -5.30885 | 7.01668 | 617.27 | 0.728 |
| gener18S:sexF | 0.34107 | -2.50415 | 2.66242 | 1000 | 0.814 |
| geneV1aR:sexF | -374.95525 | -727.11558 | -2.64809 | 1.614 | <0.001 |
| geneD1R:socialSol | -1.32173 | -4.372 | 1.27787 | 1223.358 | 0.324 |
| geneD2R:socialSol | 0.73425 | -1.64315 | 2.94601 | 929.199 | 0.554 |
| geneITR:socialSol | 0.26318 | -2.42858 | 3.1579 | 893.813 | 0.872 |
| geneMOR:socialSol | 1.66246 | -4.39772 | 7.06733 | 1000 | 0.46 |

| | | | | | |
|---|---|---|---|---|---|
| gener18S:socialSol | 0.14103 | -2.14284 | 2.68265 | 867.14 | 0.912 |
| geneV1aR:socialSol | -0.41867 | -6.27297 | 5.60197 | 763.458 | 0.862 |
| geneD1R:sexF:socialSol | 0.9841 | -2.6053 | 4.49957 | 1000 | 0.566 |
| geneD2R:sexF:socialSol | -1.66887 | -4.9295 | 1.42726 | 1000 | 0.282 |
| geneITR:sexF:socialSol | 1.29786 | -2.33542 | 5.18657 | 878.854 | 0.49 |
| geneMOR:sexF:socialSol | 1.19054 | -7.03369 | 9.27663 | 638.672 | 0.766 |
| gener18S:sexF:socialSol | 0.43835 | -3.4251 | 3.29374 | 1000 | 0.802 |
| geneV1aR:sexF:socialSol | 375.32016 | 4.42487 | 728.01143 | 1.721 | <0.001 |

**Vc (Str.CP) Pair-wise p values**

| | | | | | |
|---|---|---|---|---|---|
| $D1R | | | | | |
| | difference | | | | |
| pvalue | | sexM:socialPB | sexM:socialSol | sexF:socialPB | sexF:socialSol |
| | sexM:socialPB | NA | -1.90685 | -0.41353 | -0.90062 |
| | sexM:socialSol | 0.351856 | NA | 1.4933177 | 1.00623 |
| | sexF:socialPB | 0.840991 | 0.430851 | NA | -0.48709 |
| | sexF:socialSol | 0.639282 | 0.54399 | 0.7721882 | NA |
| $D2R | | | | | |
| | difference | | | | |
| pvalue | | sexM:socialPB | sexM:socialSol | sexF:socialPB | sexF:socialSol |
| | sexM:socialPB | NA | 1.059301 | 1.9049018 | 0.556533 |
| | sexM:socialSol | 0.543706 | NA | 0.8456005 | -0.50277 |
| | sexF:socialPB | 0.279922 | 0.641229 | NA | -1.34837 |
| | sexF:socialSol | 0.733316 | 0.755973 | 0.3928179 | NA |
| $ITR | | | | | |
| | difference | | | | |
| pvalue | | sexM:socialPB | sexM:socialSol | sexF:socialPB | sexF:socialSol |
| | sexM:socialPB | NA | 0.37969 | -0.9537568 | 1.298349 |
| | sexM:socialSol | 0.858353 | NA | -1.3334468 | 0.918659 |
| | sexF:socialPB | 0.647843 | 0.535344 | NA | 2.252106 |
| | sexF:socialSol | 0.48605 | 0.617881 | 0.232533 | NA |
| $MOR | | | | | |
| | difference | | | | |
| pvalue | | sexM:socialPB | sexM:socialSol | sexF:socialPB | sexF:socialSol |
| | sexM:socialPB | NA | 2.39842 | 1.2179173 | 5.333917 |
| | sexM:socialSol | 0.553849 | NA | -1.1805031 | 2.935497 |
| | sexF:socialPB | 0.780032 | 0.779456 | NA | 4.116 |
| | sexF:socialSol | 0.155075 | 0.439936 | 0.3113202 | NA |
| $r18S | | | | | |
| | difference | | | | |
| pvalue | | sexM:socialPB | sexM:socialSol | sexF:socialPB | sexF:socialSol |
| | sexM:socialPB | NA | 0.203459 | 0.4920597 | 1.32792 |

| | | | | | |
|---|---|---|---|---|---|
| | sexM:socialSol | 0.910904 | NA | 0.2886004 | 1.124461 |
| | sexF:socialPB | 0.793376 | 0.878347 | NA | 0.83586 |
| | sexF:socialSol | 0.431022 | 0.500693 | 0.6291014 | NA |
| $V1aR | | | | | |
| | difference | | | | |
| pvalue | | sexM:socialPB | sexM:socialSol | sexF:socialPB | sexF:socialSol |
| | sexM:socialPB | NA | -0.60402 | -540.94607 | -0.07755 |
| | sexM:socialSol | 0.915685 | NA | -540.34206 | 0.526464 |
| | sexF:socialPB | 0.10121 | 0.101881 | NA | 540.8685 |
| | sexF:socialSol | 0.989267 | 0.924626 | 0.1012361 | NA |

| **TPp (VTA)** | post.mean | l-95% | u-95% | eff.samp | pMCMC |
|---|---|---|---|---|---|
| geneD1R | 2.04395 | 0.19165 | 4.39896 | 1000 | 0.054 |
| geneD2R | 2.74824 | 0.98103 | 4.55404 | 600.48 | 0.006 |
| geneITR | -1.82547 | -3.92205 | 0.5342 | 1000 | 0.092 |
| geneMOR | 1.52105 | -0.27599 | 3.50026 | 1000 | 0.114 |
| gener18S | 12.27213 | 10.38986 | 13.76381 | 1000 | <0.001 |
| geneV1aR | -6.93717 | -19.51797 | 0.31516 | 66.53 | 0.016 |
| geneD1R:sexF | 1.83103 | -1.22826 | 4.87981 | 1000 | 0.252 |
| geneD2R:sexF | -0.01939 | -2.60002 | 2.54214 | 1000 | 0.988 |
| geneITR:sexF | 0.50735 | -2.68307 | 3.5884 | 1000 | 0.75 |
| geneMOR:sexF | -0.16825 | -2.85415 | 2.48163 | 1328.47 | 0.922 |
| gener18S:sexF | -0.496 | -2.62555 | 2.01648 | 1051.41 | 0.714 |
| geneV1aR:sexF | 3.63786 | -6.45656 | 14.81306 | 350.27 | 0.306 |
| geneD1R:socialSol | -0.47507 | -3.76198 | 2.72547 | 992.33 | 0.772 |
| geneD2R:socialSol | 0.26083 | -2.5545 | 3.64153 | 884.18 | 0.886 |
| geneITR:socialSol | 0.79589 | -2.78697 | 3.75057 | 1000 | 0.638 |
| geneMOR:socialSol | 1.03883 | -2.15066 | 4.12994 | 1202.87 | 0.544 |
| gener18S:socialSol | 0.90583 | -1.96286 | 3.70162 | 1000 | 0.514 |
| geneV1aR:socialSol | 3.16256 | -7.2433 | 15.28961 | 263.38 | 0.428 |
| geneD1R:sexF:socialSol | -2.76729 | -6.94053 | 1.54022 | 1000 | 0.18 |
| geneD2R:sexF:socialSol | -0.42801 | -4.29555 | 3.77439 | 973.19 | 0.832 |
| geneITR:sexF:socialSol | -0.56864 | -5.02669 | 3.60872 | 1000 | 0.794 |
| geneMOR:sexF:socialSol | -1.15963 | -5.55014 | 3.05552 | 1000 | 0.602 |
| gener18S:sexF:socialSol | 0.15373 | -3.27617 | 4.00776 | 1000 | 0.95 |
| geneV1aR:sexF:socialSol | -1.28967 | -14.42093 | 15.54396 | 1009.13 | 0.798 |

**TPp (VTA) Pair-wise p values**

| | | | | | |
|---|---|---|---|---|---|
| $D1R | | | | | |
| | difference | | | | |
| pvalue | | sexM:socialPB | sexM:socialSol | sexF:socialPB | sexF:socialSol |
| | sexM:socialPB | NA | -0.68537 | 2.64162402 | -2.03611 |
| | sexM:socialSol | 0.764073 | NA | 3.32699799 | -1.35073 |
| | sexF:socialPB | 0.249946 | 0.174 | NA | -4.67773 |

| | | | | | |
|---|---|---|---|---|---|
| | sexF:socialSol | 0.341659 | 0.543793 | 0.03582548 | NA |
| $D2R | | | | | |
| | difference | | | | |
| pvalue | | sexM:socialPB | sexM:socialSol | sexF:socialPB | sexF:socialSol |
| | sexM:socialPB | NA | 0.376301 | -0.0279669 | -0.26916 |
| | sexM:socialSol | 0.867176 | NA | -0.4042675 | -0.64546 |
| | sexF:socialPB | 0.988275 | 0.860694 | NA | -0.24119 |
| | sexF:socialSol | 0.886955 | 0.776691 | 0.897646 | NA |
| $ITR | | | | | |
| | difference | | | | |
| pvalue | | sexM:socialPB | sexM:socialSol | sexF:socialPB | sexF:socialSol |
| | sexM:socialPB | NA | 1.148229 | 0.7319514 | 1.059805 |
| | sexM:socialSol | 0.630556 | NA | -0.4162777 | -0.08842 |
| | sexF:socialPB | 0.752724 | 0.85878 | NA | 0.327854 |
| | sexF:socialSol | 0.628487 | 0.968866 | 0.8778985 | NA |
| $MOR | | | | | |
| | difference | | | | |
| pvalue | | sexM:socialPB | sexM:socialSol | sexF:socialPB | sexF:socialSol |
| | sexM:socialPB | NA | 1.49871 | -0.2427319 | -0.41702 |
| | sexM:socialSol | 0.52137 | NA | -1.741442 | -1.91573 |
| | sexF:socialPB | 0.904496 | 0.454887 | NA | -0.17429 |
| | sexF:socialSol | 0.83267 | 0.412547 | 0.9343204 | NA |
| $r18S | | | | | |
| | difference | | | | |
| pvalue | | sexM:socialPB | sexM:socialSol | sexF:socialPB | sexF:socialSol |
| | sexM:socialPB | NA | 1.306836 | -0.7155704 | 0.813057 |
| | sexM:socialSol | 0.527847 | NA | -2.0224059 | -0.49378 |
| | sexF:socialPB | 0.687863 | 0.345876 | NA | 1.528628 |
| | sexF:socialSol | 0.654031 | 0.814344 | 0.3975562 | NA |
| $V1aR | | | | | |
| | difference | | | | |
| pvalue | | sexM:socialPB | sexM:socialSol | sexF:socialPB | sexF:socialSol |
| | sexM:socialPB | NA | 4.562612 | 5.248318 | 7.950331 |
| | sexM:socialSol | 0.591138 | NA | 0.6857063 | 3.38772 |
| | sexF:socialPB | 0.518361 | 0.924113 | NA | 2.702013 |
| | sexF:socialSol | 0.343416 | 0.640093 | 0.6837805 | NA |

*Key*: post. mean = posterior mean, l-95% = lower 95% credible interval limit, u-95%= upper 95% credible interval limit, eff.samp = effective sample size, pMCMC = Bayesian two-tailed p-value at alpha = 0.05, *Brain region abbreviations:* teleost: telen. = telencephalon, Dm = medial part of the dorsal telen., Vd = dorsal part of the ventral telen., Dl = lateral part of the dorsal telen., Vv/Vl = lateral and ventral part of the ventral telen., Vs = supracommissural part of the ventral telen., Vc = central part of the ventral telen., POA = pre optic area, TPp = periventricular part of the posterior tuberculum., putative mammalian homolog: blAMY = basolateral amygdala, NAcc = nucleus accumbens, HIP = hippocampus, LS = lateral septum, meAMY/BNST = medial amygdala/bed nucleus of the stria terminalis, Str = Striatum, CP = caudate putamen, VTA = ventral tegmental area.

**Supplementary Table S4:** Model summary of main and interactive effects of sex and social system on brain region-specific gene expression differences in *Chaetodon* butterflyfish species. Results are reported as natural log-fold changes from the *a priori* comparison state of male pair bonding, with a two-tailed p value. Associated pair-wise differences between treatments (upper triangle) and their statistical significance (lower triangle) are reported for each brain region and gene, below summary statistics.

| **Dm (blAMY)** | post.mean | l-95% | u-95% | eff.samp | pMCMC |
|---|---|---|---|---|---|
| geneD1R | 1.44574 | 0.22494 | 2.68997 | 1000 | 0.038 |
| geneD2R | 2.62249 | 1.81163 | 3.44395 | 1000 | <0.001 |
| geneITR | -3.10927 | -4.28152 | -1.80564 | 1000 | <0.001 |
| geneMOR | 1.70223 | 0.58824 | 2.83608 | 1000 | 0.006 |
| gener18S | 13.53278 | 12.80904 | 14.14786 | 1000 | <0.001 |
| geneV1aR | -1.05394 | -1.96855 | -0.09502 | 1000 | 0.022 |
| geneD1R:sexF | -0.14586 | -1.88106 | 1.66529 | 1000 | 0.87 |
| geneD2R:sexF | -0.28945 | -1.42643 | 0.96102 | 817.5 | 0.618 |
| geneITR:sexF | -0.02948 | -1.84844 | 1.62388 | 1000 | 0.944 |
| geneMOR:sexF | 0.0985 | -1.55249 | 1.61293 | 1000 | 0.91 |
| gener18S:sexF | 0.0779 | -0.76861 | 0.84981 | 1000 | 0.852 |
| geneV1aR:sexF | 1.00629 | -0.13455 | 2.38914 | 1000 | 0.108 |
| geneD1R:socialSol | 2.23853 | -0.914 | 5.52428 | 1000 | 0.13 |
| geneD2R:socialSol | -0.57246 | -2.71433 | 1.72687 | 1000 | 0.622 |
| geneITR:socialSol | 2.13715 | -0.41862 | 4.80307 | 941.7 | 0.1 |
| geneMOR:socialSol | 1.36213 | -1.71427 | 4.11056 | 1000 | 0.35 |
| gener18S:socialSol | 0.47527 | -0.38195 | 1.28566 | 1242.6 | 0.27 |
| geneV1aR:socialSol | -2.91336 | -6.60431 | 0.63693 | 1000 | 0.088 |
| geneD1R:sexF:socialSol | -0.44821 | -4.06032 | 2.83105 | 790.7 | 0.8 |
| geneD2R:sexF:socialSol | 0.73804 | -1.9297 | 2.91302 | 1000 | 0.56 |
| geneITR:sexF:socialSol | -2.90295 | -6.06224 | 0.42407 | 808.7 | 0.064 |
| geneMOR:sexF:socialSol | -1.03257 | -4.24476 | 2.07272 | 909.7 | 0.522 |
| gener18S:sexF:socialSol | 0.36122 | -0.42896 | 1.26745 | 1000 | 0.426 |
| geneV1aR:sexF:socialSol | 0.79212 | -3.12559 | 4.80783 | 1000 | 0.718 |

**Dm (blAMY) Pair-wise p values**

| | | | | | |
|---|---|---|---|---|---|
| $D1R | | | | | |
| | difference | | | | |
| pvalue | | sexM:socialPB | sexM:socialSol | sexF:socialPB | sexF:socialSol |
| | sexM:socialPB | NA | 3.22952 | -0.21043 | 2.372465 |
| | sexM:socialSol | 0.154205 | NA | -3.43995 | -0.85706 |
| | sexF:socialPB | 0.873949 | 0.153178 | NA | 2.58289 |
| | sexF:socialSol | 0.036305 | 0.700799 | 0.02824 | NA |
| $D2R | | | | | |
| | difference | | | | |

| | | | | | |
|---|---|---|---|---|---|
| pvalue | | sexM:socialPB | sexM:socialSol | sexF:socialPB | sexF:socialSol |
| | sexM:socialPB | NA | -0.82588 | -0.41759 | -0.17871 |
| | sexM:socialSol | 0.613352 | NA | 0.408292 | 0.647172 |
| | sexF:socialPB | 0.632868 | 0.806075 | NA | 0.23888 |
| | sexF:socialSol | 0.822765 | 0.686071 | 0.761563 | NA |
| $ITR | | | | | |
| | difference | | | | |
| pvalue | | sexM:socialPB | sexM:socialSol | sexF:socialPB | sexF:socialSol |
| | sexM:socialPB | NA | 3.083252 | -0.04253 | -1.14735 |
| | sexM:socialSol | 0.107639 | NA | -3.12578 | -4.23061 |
| | sexF:socialPB | 0.973035 | 0.102134 | NA | -1.10482 |
| | sexF:socialSol | 0.354609 | 0.029861 | 0.365888 | NA |
| $MOR | | | | | |
| | difference | | | | |
| pvalue | | sexM:socialPB | sexM:socialSol | sexF:socialPB | sexF:socialSol |
| | sexM:socialPB | NA | 1.965144 | 0.142103 | 0.617563 |
| | sexM:socialSol | 0.359579 | NA | -1.82304 | -1.34758 |
| | sexF:socialPB | 0.902547 | 0.391523 | NA | 0.475459 |
| | sexF:socialSol | 0.556523 | 0.506906 | 0.64999 | NA |
| $r18S | | | | | |
| | difference | | | | |
| pvalue | | sexM:socialPB | sexM:socialSol | sexF:socialPB | sexF:socialSol |
| | sexM:socialPB | NA | 0.685669 | 0.112393 | 1.319196 |
| | sexM:socialSol | 0.266585 | NA | -0.57328 | 0.633528 |
| | sexF:socialPB | 0.846544 | 0.522219 | NA | 1.206804 |
| | sexF:socialSol | 0.044211 | 0.396899 | 0.046865 | NA |
| $V1aR | | | | | |
| | difference | | | | |
| pvalue | | sexM:socialPB | sexM:socialSol | sexF:socialPB | sexF:socialSol |
| | sexM:socialPB | NA | -4.20309 | 1.451776 | -1.60853 |
| | sexM:socialSol | 0.120428 | NA | 5.654864 | 2.594563 |
| | sexF:socialPB | 0.12302 | 0.032155 | NA | -3.0603 |
| | sexF:socialSol | 0.079061 | 0.331076 | 0.000561 | NA |

| **DI (Hip)** | post.mean | l-95% | u-95% | eff.samp | pMCMC |
|---|---|---|---|---|---|
| geneD1R | 2.83603 | 2.00926 | 3.65276 | 1000 | <0.001 |
| geneD2R | 3.88142 | 3.03286 | 4.7382 | 1000 | <0.001 |
| geneITR | -1.32316 | -2.23486 | -0.3028 | 726.7 | 0.012 |
| geneMOR | 2.77672 | 1.94175 | 3.55915 | 1000 | <0.001 |
| gener18S | 13.60635 | 12.95256 | 14.25684 | 1000 | <0.001 |
| geneV1aR | -1.62401 | -2.58845 | -0.5703 | 1000 | <0.001 |
| geneD1R:sexF | -0.54235 | -1.76821 | 0.54665 | 1000 | 0.378 |
| geneD2R:sexF | -0.31623 | -1.41107 | 0.93808 | 1000 | 0.574 |
| geneITR:sexF | 0.48435 | -0.91787 | 1.65968 | 1000 | 0.448 |
| geneMOR:sexF | -0.45118 | -1.49802 | 0.68995 | 1000 | 0.416 |
| gener18S:sexF | 0.02251 | -0.67669 | 0.8399 | 1000 | 0.976 |
| geneV1aR:sexF | 0.1891 | -1.31632 | 1.48655 | 1000 | 0.794 |
| geneD1R:socialSol | -2.22454 | -4.0422 | -0.19688 | 1000 | 0.014 |
| geneD2R:socialSol | -1.12374 | -3.29766 | 0.73992 | 1000 | 0.284 |
| geneITR:socialSol | -0.8278 | -3.11753 | 1.62734 | 1000 | 0.486 |
| geneMOR:socialSol | -0.40694 | -2.23843 | 1.65738 | 717.5 | 0.672 |
| gener18S:socialSol | 0.61907 | -0.21653 | 1.396 | 1000 | 0.152 |
| geneV1aR:socialSol | -1.08298 | -3.68386 | 1.43255 | 1000 | 0.43 |
| geneD1R:sexF:socialSol | 3.00343 | 0.84556 | 5.27253 | 1118.6 | 0.01 |
| geneD2R:sexF:socialSol | 1.21415 | -1.08482 | 3.64262 | 1000 | 0.314 |
| geneITR:sexF:socialSol | 0.97589 | -1.76516 | 3.73528 | 1000 | 0.474 |
| geneMOR:sexF:socialSol | 0.89322 | -1.29719 | 2.98213 | 891.2 | 0.424 |
| gener18S:sexF:socialSol | 0.65615 | -0.29263 | 1.47405 | 867.5 | 0.166 |
| geneV1aR:sexF:socialSol | 1.50322 | -1.33647 | 4.45025 | 1000 | 0.316 |

**DI (Hip) Pair-wise p values**

| | | | | | |
|---|---|---|---|---|---|
| $D1R | | | | | |
| | difference | | | | |
| pvalue | | sexM:socialPB | sexM:socialSol | sexF:socialPB | sexF:socialSol |
| | sexM:socialPB | NA | -3.20933 | -0.78244 | 0.341253 |
| | sexM:socialSol | 0.026028 | NA | 2.426885 | 3.550583 |
| | sexF:socialPB | 0.37126 | 0.102441 | NA | 1.123698 |
| | sexF:socialSol | 0.670096 | 0.012692 | 0.158273 | NA |
| $D2R | | | | | |
| | difference | | | | |
| pvalue | | sexM:socialPB | sexM:socialSol | sexF:socialPB | sexF:socialSol |
| | sexM:socialPB | NA | -1.62122 | -0.45622 | -0.32579 |
| | sexM:socialSol | 0.289302 | NA | 1.164997 | 1.29543 |
| | sexF:socialPB | 0.60902 | 0.444812 | NA | 0.130432 |
| | sexF:socialSol | 0.694837 | 0.396618 | 0.87223 | NA |
| $ITR | | | | | |
| | difference | | | | |

| | | | | | |
|---|---|---|---|---|---|
| pvalue | | sexM:socialPB | sexM:socialSol | sexF:socialPB | sexF:socialSol |
| | sexM:socialPB | NA | -1.19426 | 0.698769 | 0.912419 |
| | sexM:socialSol | 0.508303 | NA | 1.89303 | 2.10668 |
| | sexF:socialPB | 0.461424 | 0.298583 | NA | 0.21365 |
| | sexF:socialSol | 0.309439 | 0.242602 | 0.806035 | NA |
| $MOR | | | | | |
| | difference | | | | |
| pvalue | | sexM:socialPB | sexM:socialSol | sexF:socialPB | sexF:socialSol |
| | sexM:socialPB | NA | -0.58709 | -0.65091 | 0.050645 |
| | sexM:socialSol | 0.685328 | NA | -0.06382 | 0.637739 |
| | sexF:socialPB | 0.412403 | 0.964847 | NA | 0.701555 |
| | sexF:socialSol | 0.948753 | 0.654671 | 0.360568 | NA |
| $r18S | | | | | |
| | difference | | | | |
| pvalue | | sexM:socialPB | sexM:socialSol | sexF:socialPB | sexF:socialSol |
| | sexM:socialPB | NA | 0.893134 | 0.032469 | 1.872228 |
| | sexM:socialSol | 0.139886 | NA | -0.86067 | 0.979094 |
| | sexF:socialPB | 0.953443 | 0.290072 | NA | 1.83976 |
| | sexF:socialSol | 0.005993 | 0.17752 | 0.002714 | NA |
| $V1aR | | | | | |
| | difference | | | | |
| pvalue | | sexM:socialPB | sexM:socialSol | sexF:socialPB | sexF:socialSol |
| | sexM:socialPB | NA | -1.56241 | 0.272819 | 0.879098 |
| | sexM:socialSol | 0.422236 | NA | 1.835228 | 2.441507 |
| | sexF:socialPB | 0.79077 | 0.347525 | NA | 0.606279 |
| | sexF:socialSol | 0.328776 | 0.201258 | 0.514288 | NA |

| **Vv/vl (LS)** | post.mean | l-95% | u-95% | eff.samp | pMCMC |
|---|---|---|---|---|---|
| geneD1R | 3.5832 | 2.489433 | 4.775977 | 1000 | <0.001 |
| geneD2R | 5.298092 | 4.186972 | 6.382995 | 1000 | <0.001 |
| geneITR | 0.609692 | -0.59512 | 1.670394 | 1764.69 | 0.314 |
| geneMOR | 4.973526 | 3.787024 | 6.112364 | 1000 | <0.001 |
| gener18S | 13.55028 | 12.44062 | 14.57589 | 713.78 | <0.001 |
| geneV1aR | -0.51312 | -1.78853 | 0.615867 | 1000 | 0.41 |
| geneD1R:sexF | -0.73147 | -2.48378 | 0.909196 | 907.46 | 0.382 |
| geneD2R:sexF | -0.74199 | -2.4489 | 0.894267 | 1608.73 | 0.37 |
| geneITR:sexF | -1.79562 | -3.86023 | -0.05581 | 1000 | 0.06 |
| geneMOR:sexF | -1.60171 | -3.40637 | 0.194791 | 843.32 | 0.074 |
| gener18S:sexF | -1.11077 | -2.77129 | 0.590953 | 893.51 | 0.184 |
| geneV1aR:sexF | -0.00968 | -2.03937 | 1.677377 | 864.35 | 0.99 |
| geneD1R:socialSol | -0.98122 | -4.56126 | 2.151644 | 1000 | 0.568 |
| geneD2R:socialSol | -1.60931 | -4.97809 | 0.845963 | 1000 | 0.248 |
| geneITR:socialSol | -21.8665 | -50.8821 | -2.58244 | 18.94 | 0.006 |
| geneMOR:socialSol | -2.22592 | -4.99823 | 0.953448 | 988.09 | 0.144 |

| | | | | | |
|---|---|---|---|---|---|
| gener18S:socialSol | 0.405981 | -2.47623 | 3.58375 | 1000 | 0.8 |
| geneV1aR:socialSol | -0.02057 | -3.00152 | 3.337803 | 780.22 | 0.962 |
| geneD1R:sexF:socialSol | 0.941765 | -2.72746 | 4.455502 | 911.94 | 0.61 |
| geneD2R:sexF:socialSol | 0.110958 | -2.98804 | 3.564392 | 1000 | 0.944 |
| geneITR:sexF:socialSol | 21.1941 | 1.530546 | 50.45012 | 18.98 | 0.01 |
| geneMOR:sexF:socialSol | 1.219434 | -2.39012 | 4.438878 | 1000 | 0.488 |
| gener18S:sexF:socialSol | 0.617217 | -2.84638 | 4.008031 | 1000 | 0.72 |
| geneV1aR:sexF:socialSol | -0.71249 | -4.06526 | 3.532017 | 822.51 | 0.734 |

**Vv/vl (LS) Pair-wise p values**

| | | | | | |
|---|---|---|---|---|---|
| $D1R | | | | | |
| | difference | | | | |
| pvalue | | sexM:socialPB | sexM:socialSol | sexF:socialPB | sexF:socialSol |
| | sexM:socialPB | NA | -1.11199 | -1.00306 | -1.12236 |
| | sexM:socialSol | 0.652415 | NA | 0.108926 | -0.01038 |
| | sexF:socialPB | 0.433234 | 0.965808 | NA | -0.1193 |
| | sexF:socialSol | 0.310196 | 0.99668 | 0.915758 | NA |
| $D2R | | | | | |
| | difference | | | | |
| pvalue | | sexM:socialPB | sexM:socialSol | sexF:socialPB | sexF:socialSol |
| | sexM:socialPB | NA | -2.38907 | -1.10505 | -3.26143 |
| | sexM:socialSol | 0.275636 | NA | 1.284018 | -0.87236 |
| | sexF:socialPB | 0.380425 | 0.581213 | NA | -2.15637 |
| | sexF:socialSol | 0.002987 | 0.694601 | 0.060225 | NA |
| $ITR | | | | | |
| | difference | | | | |
| pvalue | | sexM:socialPB | sexM:socialSol | sexF:socialPB | sexF:socialSol |
| | sexM:socialPB | NA | -69.5825 | -2.69084 | -3.49217 |
| | sexM:socialSol | 0.03968 | NA | 66.89166 | 66.09033 |
| | sexF:socialPB | 0.057969 | 0.048714 | NA | -0.80133 |
| | sexF:socialSol | 0.002544 | 0.050417 | 0.566333 | NA |
| $MOR | | | | | |
| | difference | | | | |
| pvalue | | sexM:socialPB | sexM:socialSol | sexF:socialPB | sexF:socialSol |
| | sexM:socialPB | NA | -3.25798 | -2.38402 | -3.78774 |
| | sexM:socialSol | 0.125114 | NA | 0.87396 | -0.52976 |
| | sexF:socialPB | 0.071924 | 0.69984 | NA | -1.40372 |
| | sexF:socialSol | 0.001037 | 0.805827 | 0.288721 | NA |
| $r18S | | | | | |
| | difference | | | | |
| pvalue | | sexM:socialPB | sexM:socialSol | sexF:socialPB | sexF:socialSol |
| | sexM:socialPB | NA | 0.361916 | -1.74085 | -0.18854 |
| | sexM:socialSol | 0.861812 | NA | -2.10276 | -0.55045 |

| | | | | | |
|---|---|---|---|---|---|
| | sexF:socialPB | 0.17367 | 0.340323 | NA | 1.552312 |
| | sexF:socialSol | 0.863452 | 0.793924 | 0.212397 | NA |
| $V1aR | | | | | |
| | difference | | | | |
| pvalue | | sexM:socialPB | sexM:socialSol | sexF:socialPB | sexF:socialSol |
| | sexM:socialPB | NA | -0.01282 | 0.086331 | -0.97882 |
| | sexM:socialSol | 0.995491 | NA | 0.099154 | -0.966 |
| | sexF:socialPB | 0.950805 | 0.9667 | NA | -1.06516 |
| | sexF:socialSol | 0.41617 | 0.673975 | 0.416138 | NA |

| **Vs (meAMY/BNST)** | post.mean | l-95% | u-95% | eff.samp | pMCMC |
|---|---|---|---|---|---|
| geneD1R | -0.02012 | -1.58107 | 1.27648 | 1000 | 0.986 |
| geneD2R | 2.7909 | 1.7242 | 3.72432 | 1000 | <0.001 |
| geneITR | -1.78011 | -2.90743 | -0.63882 | 1000 | <0.001 |
| geneMOR | 3.82531 | 2.21181 | 5.19858 | 1000 | <0.001 |
| gener18S | 12.93983 | 12.22084 | 13.65862 | 1000 | <0.001 |
| geneV1aR | -0.20625 | -1.17062 | 0.81149 | 1000 | 0.706 |
| geneD1R:sexF | 0.48622 | -1.2947 | 2.55869 | 1280.198 | 0.622 |
| geneD2R:sexF | -1.07103 | -2.5067 | 0.57119 | 1000 | 0.17 |
| geneITR:sexF | -0.34163 | -2.0711 | 1.24912 | 1000 | 0.688 |
| geneMOR:sexF | -1.39005 | -3.42641 | 0.63305 | 890.407 | 0.174 |
| gener18S:sexF | -0.14552 | -1.00938 | 0.6635 | 1000 | 0.74 |
| geneV1aR:sexF | -1.20378 | -2.65877 | 0.29088 | 1134.831 | 0.116 |
| geneD1R:socialSol | 2.28408 | -0.82678 | 4.89858 | 1000 | 0.122 |
| geneD2R:socialSol | -0.80521 | -3.36659 | 1.63126 | 1000 | 0.546 |
| geneITR:socialSol | -62.7912 | -107.403 | -10.6826 | 6.166 | <0.001 |
| geneMOR:socialSol | -2.142 | -5.12538 | 1.26886 | 825.33 | 0.19 |
| gener18S:socialSol | 0.25603 | -0.57557 | 1.12704 | 1044.283 | 0.58 |
| geneV1aR:socialSol | -2.48793 | -5.46044 | 0.27616 | 1110.833 | 0.094 |
| geneD1R:sexF:socialSol | -1.69031 | -4.90092 | 1.47637 | 1141.286 | 0.324 |
| geneD2R:sexF:socialSol | 0.50767 | -2.18516 | 3.55699 | 1000 | 0.732 |
| geneITR:sexF:socialSol | 61.62759 | 9.03708 | 106.7488 | 6.172 | 0.002 |
| geneMOR:sexF:socialSol | 1.68438 | -2.25177 | 5.23685 | 951.973 | 0.352 |
| gener18S:sexF:socialSol | 0.4844 | -0.47065 | 1.47312 | 1000 | 0.266 |
| geneV1aR:sexF:socialSol | 0.39478 | -2.4252 | 4.16885 | 1145.064 | 0.826 |

**Vs (meAMY/BNST) Pair-wise p values**

| | | | | | |
|---|---|---|---|---|---|
| $D1R | | | | | |
| | difference | | | | |
| pvalue | | sexM:socialPB | sexM:socialSol | sexF:socialPB | sexF:socialSol |
| | sexM:socialPB | NA | 3.295237 | 0.70147 | 1.5581 |
| | sexM:socialSol | 0.121654 | NA | -2.59377 | -1.73714 |
| | sexF:socialPB | 0.630149 | 0.254291 | NA | 0.85663 |
| | sexF:socialSol | 0.216014 | 0.393439 | 0.514274 | NA |

| | | | | | |
|---|---|---|---|---|---|
| $D2R | | | | | |
| | difference | | | | |
| pvalue | | sexM:socialPB | sexM:socialSol | sexF:socialPB | sexF:socialSol |
| | sexM:socialPB | NA | -1.16167 | -1.54517 | -1.97443 |
| | sexM:socialSol | 0.537387 | NA | -0.3835 | -0.81276 |
| | sexF:socialPB | 0.170544 | 0.842261 | NA | -0.42926 |
| | sexF:socialSol | 0.040983 | 0.659846 | 0.669985 | NA |
| $ITR | | | | | |
| | difference | | | | |
| pvalue | | sexM:socialPB | sexM:socialSol | sexF:socialPB | sexF:socialSol |
| | sexM:socialPB | NA | -90.5886 | -0.49287 | -2.17165 |
| | sexM:socialSol | 0.017358 | NA | 90.09572 | 88.41694 |
| | sexF:socialPB | 0.689377 | 0.017821 | NA | -1.67878 |
| | sexF:socialSol | 0.064221 | 0.020139 | 0.157218 | NA |
| $MOR | | | | | |
| | difference | | | | |
| pvalue | | sexM:socialPB | sexM:socialSol | sexF:socialPB | sexF:socialSol |
| | sexM:socialPB | NA | -3.09025 | -2.00542 | -2.66562 |
| | sexM:socialSol | 0.199156 | NA | 1.084839 | 0.424631 |
| | sexF:socialPB | 0.189659 | 0.655901 | NA | -0.66021 |
| | sexF:socialSol | 0.052704 | 0.852147 | 0.625882 | NA |
| $r18S | | | | | |
| | difference | | | | |
| pvalue | | sexM:socialPB | sexM:socialSol | sexF:socialPB | sexF:socialSol |
| | sexM:socialPB | NA | 0.369375 | -0.20994 | 0.858281 |
| | sexM:socialSol | 0.57556 | NA | -0.57931 | 0.488907 |
| | sexF:socialPB | 0.734772 | 0.535833 | NA | 1.068219 |
| | sexF:socialSol | 0.231286 | 0.515069 | 0.118097 | NA |
| $V1aR | | | | | |
| | difference | | | | |
| pvalue | | sexM:socialPB | sexM:socialSol | sexF:socialPB | sexF:socialSol |
| | sexM:socialPB | NA | -3.58932 | -1.73669 | -4.75646 |
| | sexM:socialSol | 0.095338 | NA | 1.852629 | -1.16714 |
| | sexF:socialPB | 0.111333 | 0.389553 | NA | -3.01977 |
| | sexF:socialSol | 3.8E-05 | 0.592622 | 0.008905 | NA |

| **Vd (NAcc)** | post.mean | l-95% | u-95% | eff.samp | pMCMC |
|---|---|---|---|---|---|
| geneD1R | 0.030583 | -1.05863 | 1.195641 | 858.5 | 0.934 |
| geneD2R | 1.045259 | 0.103456 | 2.026887 | 1000 | 0.044 |
| geneITR | -6.38492 | -9.32547 | -3.45683 | 453.3 | <0.001 |
| geneMOR | -0.37873 | -2.15279 | 1.173196 | 1000 | 0.7 |
| gener18S | 11.49026 | 10.80712 | 12.08588 | 1000 | <0.001 |
| geneV1aR | -3.44397 | -4.94704 | -2.00824 | 1000 | <0.001 |

| | | | | | |
|---|---|---|---|---|---|
| geneD1R:sexF | 1.165669 | -0.3764 | 2.755153 | 1000 | 0.154 |
| geneD2R:sexF | -0.16332 | -1.83136 | 1.149953 | 1000 | 0.846 |
| geneITR:sexF | 2.356562 | -0.48914 | 6.131001 | 599.6 | 0.118 |
| geneMOR:sexF | -0.68067 | -3.41874 | 1.952773 | 888.1 | 0.622 |
| gener18S:sexF | -0.00686 | -0.70288 | 0.791048 | 1000 | 0.986 |
| geneV1aR:sexF | 0.323769 | -1.66843 | 2.121991 | 1000 | 0.738 |
| geneD1R:socialSol | 0.474709 | -1.65255 | 2.985605 | 1000 | 0.706 |
| geneD2R:socialSol | 0.22745 | -2.31579 | 2.317808 | 1000 | 0.812 |
| geneITR:socialSol | 2.169236 | -2.24577 | 6.639113 | 539.8 | 0.338 |
| geneMOR:socialSol | 0.74441 | -2.70441 | 4.278152 | 1000 | 0.726 |
| gener18S:socialSol | 0.332802 | -0.47689 | 1.248549 | 1000 | 0.444 |
| geneV1aR:socialSol | 1.105303 | -1.67422 | 3.719492 | 1036.7 | 0.444 |
| geneD1R:sexF:socialSol | -2.02674 | -4.69551 | 0.557911 | 1000 | 0.128 |
| geneD2R:sexF:socialSol | -1.38164 | -3.90727 | 1.651573 | 1000 | 0.322 |
| geneITR:sexF:socialSol | -2.46767 | -7.37543 | 2.770347 | 703.5 | 0.336 |
| geneMOR:sexF:socialSol | 0.117548 | -4.53717 | 4.430892 | 1000 | 0.932 |
| gener18S:sexF:socialSol | 0.349958 | -0.59411 | 1.223655 | 1000 | 0.454 |
| geneV1aR:sexF:socialSol | -2.04997 | -5.42774 | 1.009508 | 995.5 | 0.21 |

**Vd (NAcc) Pair-wise p values**

| | | | | | |
|---|---|---|---|---|---|
| $D1R | | | | | |
| | difference | | | | |
| pvalue | | sexM:socialPB | sexM:socialSol | sexF:socialPB | sexF:socialSol |
| | sexM:socialPB | NA | 0.805892 | 1.706549 | -0.61999 |
| | sexM:socialSol | 0.639216 | NA | 0.900657 | -1.42588 |
| | sexF:socialPB | 0.137 | 0.606262 | NA | -2.32654 |
| | sexF:socialSol | 0.547509 | 0.388731 | 0.03047 | NA |
| $D2R | | | | | |
| | difference | | | | |
| pvalue | | sexM:socialPB | sexM:socialSol | sexF:socialPB | sexF:socialSol |
| | sexM:socialPB | NA | 0.259607 | -0.29081 | -1.90389 |
| | sexM:socialSol | 0.875115 | NA | -0.55042 | -2.16349 |
| | sexF:socialPB | 0.798517 | 0.754715 | NA | -1.61307 |
| | sexF:socialSol | 0.054982 | 0.188008 | 0.141152 | NA |
| $ITR | | | | | |
| | difference | | | | |
| pvalue | | sexM:socialPB | sexM:socialSol | sexF:socialPB | sexF:socialSol |
| | sexM:socialPB | NA | 3.129402 | 3.432013 | 3.089294 |
| | sexM:socialSol | 0.348683 | NA | 0.302611 | -0.04011 |
| | sexF:socialPB | 0.165197 | 0.914274 | NA | -0.34272 |
| | sexF:socialSol | 0.19753 | 0.98822 | 0.824439 | NA |
| $MOR | | | | | |
| | difference | | | | |
| pvalue | | sexM:socialPB | sexM:socialSol | sexF:socialPB | sexF:socialSol |

| | | | | | |
|---|---|---|---|---|---|
| | sexM:socialPB | NA | 0.999995 | -0.98402 | 0.330487 |
| | sexM:socialSol | 0.699819 | NA | -1.98401 | -0.66951 |
| | sexF:socialPB | 0.609285 | 0.482176 | NA | 1.314505 |
| | sexF:socialSol | 0.839055 | 0.793307 | 0.443256 | NA |
| $r18S | | | | | |
| | difference | | | | |
| pvalue | | sexM:socialPB | sexM:socialSol | sexF:socialPB | sexF:socialSol |
| | sexM:socialPB | NA | 0.498663 | 0.031447 | 1.01339 |
| | sexM:socialSol | 0.419681 | NA | -0.46722 | 0.514727 |
| | sexF:socialPB | 0.955814 | 0.599333 | NA | 0.981943 |
| | sexF:socialSol | 0.102503 | 0.459069 | 0.119823 | NA |
| $V1aR | | | | | |
| | difference | | | | |
| pvalue | | sexM:socialPB | sexM:socialSol | sexF:socialPB | sexF:socialSol |
| | sexM:socialPB | NA | 1.635519 | 0.532984 | -0.89915 |
| | sexM:socialSol | 0.434986 | NA | -1.10254 | -2.53467 |
| | sexF:socialPB | 0.705167 | 0.59971 | NA | -1.43213 |
| | sexF:socialSol | 0.464576 | 0.20936 | 0.269615 | NA |

| **POA (POA)** | post.mean | l-95% | u-95% | eff.samp | pMCMC |
|---|---|---|---|---|---|
| geneD1R | 3.47028 | 2.55003 | 4.31307 | 1000 | <0.001 |
| geneD2R | 4.34809 | 3.50166 | 5.17739 | 1000 | <0.001 |
| geneITR | 0.08649 | -0.81952 | 0.86331 | 1000 | 0.804 |
| geneMOR | 4.16535 | 3.23615 | 5.15936 | 1000 | <0.001 |
| geneV1aR | -0.35394 | -1.31339 | 0.70585 | 1000 | 0.494 |
| gener18S | 12.62434 | 11.9034 | 13.27657 | 1123 | <0.001 |
| geneD1R:sexF | -0.25467 | -1.51291 | 0.9873 | 1000 | 0.646 |
| geneD2R:sexF | -0.14761 | -1.36827 | 0.98772 | 1000 | 0.844 |
| geneITR:sexF | 0.27825 | -0.80299 | 1.61599 | 1000 | 0.65 |
| geneMOR:sexF | -0.16301 | -1.5625 | 1.10806 | 990.2 | 0.814 |
| geneV1aR:sexF | -0.48373 | -2.08242 | 0.70981 | 1000 | 0.512 |
| gener18S:sexF | 0.18975 | -0.48194 | 0.88497 | 1000 | 0.61 |
| geneD1R:socialSol | 0.61762 | -1.46324 | 2.66489 | 1000 | 0.584 |
| geneD2R:socialSol | -0.6874 | -2.7015 | 1.37032 | 1000 | 0.492 |
| geneITR:socialSol | -1.30697 | -3.55063 | 0.88706 | 1000 | 0.244 |
| geneMOR:socialSol | -0.68967 | -2.91948 | 1.47857 | 1000 | 0.552 |
| geneV1aR:socialSol | -0.8422 | -2.99982 | 1.50492 | 1000 | 0.48 |
| gener18S:socialSol | 0.75208 | -0.09433 | 1.62248 | 1000 | 0.108 |
| geneD1R:sexF:socialSol | 0.5488 | -1.82717 | 2.6809 | 1167.5 | 0.652 |
| geneD2R:sexF:socialSol | 1.20831 | -1.1817 | 3.37319 | 1000 | 0.298 |
| geneITR:sexF:socialSol | 1.50295 | -0.97492 | 3.93551 | 1000 | 0.238 |
| geneMOR:sexF:socialSol | 1.25509 | -1.24497 | 3.92943 | 1000 | 0.318 |
| geneV1aR:sexF:socialSol | 0.81277 | -1.91626 | 3.16279 | 1000 | 0.556 |
| gener18S:sexF:socialSol | 0.68544 | -0.20761 | 1.52233 | 1000 | 0.108 |

**POA (POA) Pair-wise p values**

| | | | | | |
|---|---|---|---|---|---|
| $D1R | | | | | |
| | difference | | | | |
| pvalue | | sexM:socialPB | sexM:socialSol | sexF:socialPB | sexF:socialSol |
| | sexM:socialPB | NA | 0.891042 | -0.36742 | 1.315372 |
| | sexM:socialSol | 0.561675 | NA | -1.25846 | 0.424331 |
| | sexF:socialPB | 0.686977 | 0.40222 | NA | 1.682788 |
| | sexF:socialSol | 0.112295 | 0.774482 | 0.032288 | NA |
| $D2R | | | | | |
| | difference | | | | |
| pvalue | | sexM:socialPB | sexM:socialSol | sexF:socialPB | sexF:socialSol |
| | sexM:socialPB | NA | -0.99171 | -0.21296 | 0.538564 |
| | sexM:socialSol | 0.508362 | NA | 0.778753 | 1.530273 |
| | sexF:socialPB | 0.816171 | 0.599704 | NA | 0.75152 |
| | sexF:socialSol | 0.498257 | 0.276759 | 0.35441 | NA |
| $ITR | | | | | |
| | difference | | | | |
| pvalue | | sexM:socialPB | sexM:socialSol | sexF:socialPB | sexF:socialSol |
| | sexM:socialPB | NA | -1.88556 | 0.401424 | 0.684165 |
| | sexM:socialSol | 0.248864 | NA | 2.28698 | 2.56972 |
| | sexF:socialPB | 0.65417 | 0.157506 | NA | 0.282741 |
| | sexF:socialSol | 0.403625 | 0.100053 | 0.733244 | NA |
| $MOR | | | | | |
| | difference | | | | |
| pvalue | | sexM:socialPB | sexM:socialSol | sexF:socialPB | sexF:socialSol |
| | sexM:socialPB | NA | -0.99498 | -0.23517 | 0.580553 |
| | sexM:socialSol | 0.551456 | NA | 0.759808 | 1.575536 |
| | sexF:socialPB | 0.812459 | 0.651128 | NA | 0.815728 |
| | sexF:socialSol | 0.510823 | 0.334236 | 0.379342 | NA |
| $V1aR | | | | | |
| | difference | | | | |
| pvalue | | sexM:socialPB | sexM:socialSol | sexF:socialPB | sexF:socialSol |
| | sexM:socialPB | NA | -1.21504 | -0.69787 | -0.74033 |
| | sexM:socialSol | 0.468099 | NA | 0.517168 | 0.47471 |
| | sexF:socialPB | 0.499057 | 0.757094 | NA | -0.04246 |
| | sexF:socialSol | 0.423515 | 0.768798 | 0.963275 | NA |
| $r18S | | | | | |
| | difference | | | | |
| pvalue | | sexM:socialPB | sexM:socialSol | sexF:socialPB | sexF:socialSol |
| | sexM:socialPB | NA | 1.085015 | 0.273756 | 2.34765 |
| | sexM:socialSol | 0.088925 | NA | -0.81126 | 1.262635 |
| | sexF:socialPB | 0.60507 | 0.347608 | NA | 2.073894 |

| | | | | | |
|---|---|---|---|---|---|
| | sexF:socialSol | 0.000266 | 0.084652 | 0.000437 | NA |

| **Vc (Str/CP)** | post.mean | l-95% | u-95% | eff.samp | pMCMC |
|---|---|---|---|---|---|
| geneD1R | 2.65814 | 1.79555 | 3.7582 | 1000 | <0.001 |
| geneD2R | 4.01137 | 3.06902 | 4.92022 | 1000 | <0.001 |
| geneITR | -0.13941 | -1.16918 | 0.85515 | 1000 | 0.78 |
| geneMOR | 3.19274 | 1.60444 | 4.75663 | 837.1 | <0.001 |
| geneV1aR | -2.00819 | -3.30649 | -0.87314 | 1000 | <0.001 |
| gener18S | 12.22007 | 11.47743 | 13.01413 | 1000 | <0.001 |
| geneD1R:sexF | -0.24252 | -1.72257 | 1.31244 | 1000 | 0.75 |
| geneD2R:sexF | 0.2277 | -1.28501 | 1.66822 | 1000 | 0.754 |
| geneITR:sexF | -0.7645 | -2.30366 | 0.73032 | 1025.2 | 0.35 |
| geneMOR:sexF | 0.91649 | -1.67809 | 3.07477 | 1000 | 0.454 |
| geneV1aR:sexF | -0.6868 | -2.71887 | 1.26281 | 990.3 | 0.508 |
| gener18S:sexF | 0.07285 | -0.77938 | 0.87739 | 1000 | 0.88 |
| geneD1R:socialSol | 0.5747 | -1.53936 | 2.64884 | 1000 | 0.598 |
| geneD2R:socialSol | -1.29885 | -3.36558 | 0.68517 | 1177.6 | 0.228 |
| geneITR:socialSol | -2.76981 | -5.75164 | -0.04515 | 1000 | 0.052 |
| geneMOR:socialSol | -1.27822 | -4.44769 | 2.09977 | 1000 | 0.448 |
| geneV1aR:socialSol | -2.51782 | -6.1199 | 1.26103 | 1000 | 0.192 |
| gener18S:socialSol | 0.39303 | -0.4916 | 1.22719 | 1000 | 0.386 |
| geneD1R:sexF:socialSol | -0.64881 | -3.18752 | 1.93847 | 838.6 | 0.612 |
| geneD2R:sexF:socialSol | 0.43111 | -2.257 | 2.84365 | 1000 | 0.758 |
| geneITR:sexF:socialSol | 2.74546 | -0.44973 | 6.22583 | 1000 | 0.088 |
| geneMOR:sexF:socialSol | -0.38578 | -4.4329 | 3.622 | 1000 | 0.812 |
| geneV1aR:sexF:socialSol | 3.20803 | -1.07125 | 7.52059 | 1000 | 0.114 |
| gener18S:sexF:socialSol | 0.30727 | -0.60411 | 1.24304 | 1000 | 0.52 |

**Vc (Str/CP) Pair-wise p values**

| | | | | | |
|---|---|---|---|---|---|
| $D1R | | | | | |
| | difference | | | | |
| pvalue | | sexM:socialPB | sexM:socialSol | sexF:socialPB | sexF:socialSol |
| | sexM:socialPB | NA | 0.82911 | -0.34988 | -0.4568 |
| | sexM:socialSol | 0.596079 | NA | -1.17899 | -1.28591 |
| | sexF:socialPB | 0.763587 | 0.470557 | NA | -0.10692 |
| | sexF:socialSol | 0.613972 | 0.391513 | 0.917392 | NA |
| $D2R | | | | | |
| | difference | | | | |
| pvalue | | sexM:socialPB | sexM:socialSol | sexF:socialPB | sexF:socialSol |
| | sexM:socialPB | NA | -1.87385 | 0.328499 | -0.92339 |
| | sexM:socialSol | 0.217034 | NA | 2.202347 | 0.950463 |
| | sexF:socialPB | 0.764915 | 0.157432 | NA | -1.25188 |
| | sexF:socialSol | 0.273209 | 0.523657 | 0.203337 | NA |
| $ITR | | | | | |

| | | | | | |
|---|---|---|---|---|---|
| | difference | | | | |
| pvalue | | sexM:socialPB | sexM:socialSol | sexF:socialPB | sexF:socialSol |
| | sexM:socialPB | NA | -3.996 | -1.10294 | -1.13808 |
| | sexM:socialSol | 0.061854 | NA | 2.893057 | 2.857914 |
| | sexF:socialPB | 0.341241 | 0.180948 | NA | -0.03514 |
| | sexF:socialSol | 0.240267 | 0.172433 | 0.974438 | NA |
| $MOR | | | | | |
| | difference | | | | |
| pvalue | | sexM:socialPB | sexM:socialSol | sexF:socialPB | sexF:socialSol |
| | sexM:socialPB | NA | -1.84408 | 1.32221 | -1.07844 |
| | sexM:socialSol | 0.449884 | NA | 3.166288 | 0.765641 |
| | sexF:socialPB | 0.461679 | 0.208495 | NA | -2.40065 |
| | sexF:socialSol | 0.469333 | 0.750179 | 0.143498 | NA |
| $V1aR | | | | | |
| | difference | | | | |
| pvalue | | sexM:socialPB | sexM:socialSol | sexF:socialPB | sexF:socialSol |
| | sexM:socialPB | NA | -3.63245 | -0.99084 | 0.004915 |
| | sexM:socialSol | 0.199562 | NA | 2.641602 | 3.637362 |
| | sexF:socialPB | 0.499566 | 0.370046 | NA | 0.99576 |
| | sexF:socialSol | 0.996439 | 0.195251 | 0.452305 | NA |
| $r18S | | | | | |
| | difference | | | | |
| pvalue | | sexM:socialPB | sexM:socialSol | sexF:socialPB | sexF:socialSol |
| | sexM:socialPB | NA | 0.567019 | 0.105095 | 1.115413 |
| | sexM:socialSol | 0.383312 | NA | -0.46192 | 0.548393 |
| | sexF:socialPB | 0.867387 | 0.623853 | NA | 1.010318 |
| | sexF:socialSol | 0.142133 | 0.478285 | 0.157653 | NA |

| **TPp (VTA)** | post.mean | l-95% | u-95% | eff.samp | pMCMC |
|---|---|---|---|---|---|
| geneD1R | 0.55477 | -0.56094 | 1.76715 | 1231.9 | 0.352 |
| geneD2R | 2.26394 | 1.17787 | 3.14334 | 1120.8 | <0.001 |
| geneITR | -1.17488 | -2.37169 | -0.05856 | 1000 | 0.056 |
| geneMOR | 2.00858 | 1.01206 | 3.00135 | 928.6 | <0.001 |
| geneV1aR | -2.63766 | -4.08429 | -1.38868 | 1000 | <0.001 |
| gener18S | 11.22024 | 10.45378 | 12.01931 | 1000 | <0.001 |
| geneD1R:sexF | 1.22318 | -0.52844 | 2.80597 | 1010 | 0.162 |
| geneD2R:sexF | 0.97531 | -0.41269 | 2.35845 | 1000 | 0.172 |
| geneITR:sexF | 0.5566 | -0.97961 | 2.17189 | 938.9 | 0.504 |
| geneMOR:sexF | 0.41159 | -1.02057 | 1.9225 | 997.5 | 0.582 |

| | | | | | |
|---|---|---|---|---|---|
| geneV1aR:sexF | 0.78672 | -0.95336 | 2.73478 | 1000 | 0.388 |
| gener18S:sexF | 0.29173 | -0.51454 | 1.13936 | 1000 | 0.51 |
| geneD1R:socialSol | 1.50616 | -0.83438 | 3.87766 | 1000 | 0.19 |
| geneD2R:socialSol | -0.09204 | -2.47839 | 2.0791 | 1000 | 0.914 |
| geneITR:socialSol | -0.48215 | -2.88487 | 2.41712 | 1146.4 | 0.73 |
| geneMOR:socialSol | 0.89084 | -1.3751 | 3.12954 | 1000 | 0.44 |
| geneV1aR:socialSol | 0.29866 | -2.41009 | 3.48911 | 1000 | 0.874 |
| gener18S:socialSol | 0.54636 | -0.3464 | 1.3792 | 1000 | 0.244 |
| geneD1R:sexF:socialSol | -1.68391 | -4.52297 | 0.98605 | 1000 | 0.232 |
| geneD2R:sexF:socialSol | -1.23208 | -3.88805 | 1.28271 | 1000 | 0.348 |
| geneITR:sexF:socialSol | -0.161 | -3.26474 | 2.81771 | 1097 | 0.924 |
| geneMOR:sexF:socialSol | -0.95217 | -3.92275 | 1.44154 | 1000 | 0.472 |
| geneV1aR:sexF:socialSol | -0.28906 | -3.69368 | 3.24725 | 1000 | 0.844 |
| gener18S:sexF:socialSol | 0.20439 | -0.71813 | 1.10849 | 951.6 | 0.678 |

**TPp (VTA) Pair-wise p values**

| | | | | | |
|---|---|---|---|---|---|
| $D1R | | | | | |
| | difference | | | | |
| pvalue | | sexM:socialPB | sexM:socialSol | sexF:socialPB | sexF:socialSol |
| | sexM:socialPB | NA | 2.172936 | 1.76467 | 1.508236 |
| | sexM:socialSol | 0.206875 | NA | -0.40827 | -0.6647 |
| | sexF:socialPB | 0.147852 | 0.820516 | NA | -0.25643 |
| | sexF:socialSol | 0.154253 | 0.690657 | 0.807971 | NA |
| $D2R | | | | | |
| | difference | | | | |
| pvalue | | sexM:socialPB | sexM:socialSol | sexF:socialPB | sexF:socialSol |
| | sexM:socialPB | NA | -0.13278 | 1.407081 | -0.50322 |
| | sexM:socialSol | 0.937058 | NA | 1.539863 | -0.37044 |
| | sexF:socialPB | 0.177968 | 0.36324 | NA | -1.9103 |
| | sexF:socialSol | 0.593096 | 0.822446 | 0.052901 | NA |
| $ITR | | | | | |
| | difference | | | | |
| pvalue | | sexM:socialPB | sexM:socialSol | sexF:socialPB | sexF:socialSol |
| | sexM:socialPB | NA | -0.69559 | 0.803004 | -0.12487 |
| | sexM:socialSol | 0.721516 | NA | 1.498596 | 0.570725 |
| | sexF:socialPB | 0.493903 | 0.42187 | NA | -0.92787 |
| | sexF:socialSol | 0.908185 | 0.760896 | 0.359986 | NA |
| $MOR | | | | | |
| | difference | | | | |
| pvalue | | sexM:socialPB | sexM:socialSol | sexF:socialPB | sexF:socialSol |

| | | | | | |
|---|---|---|---|---|---|
| | sexM:socialPB | NA | 1.285207 | 0.593792 | 0.505314 |
| | sexM:socialSol | 0.447889 | NA | -0.69141 | -0.77989 |
| | sexF:socialPB | 0.587829 | 0.683541 | NA | -0.08848 |
| | sexF:socialSol | 0.598355 | 0.625549 | 0.925786 | NA |
| $V1aR | | | | | |
| | difference | | | | |
| pvalue | | sexM:socialPB | sexM:socialSol | sexF:socialPB | sexF:socialSol |
| | sexM:socialPB | NA | 0.430869 | 1.134996 | 1.148842 |
| | sexM:socialSol | 0.845226 | NA | 0.704127 | 0.717973 |
| | sexF:socialPB | 0.395304 | 0.742335 | NA | 0.013846 |
| | sexF:socialSol | 0.357187 | 0.73087 | 0.990855 | NA |
| $r18S | | | | | |
| | difference | | | | |
| pvalue | | sexM:socialPB | sexM:socialSol | sexF:socialPB | sexF:socialSol |
| | sexM:socialPB | NA | 0.788225 | 0.420873 | 1.503977 |
| | sexM:socialSol | 0.225171 | NA | -0.36735 | 0.715752 |
| | sexF:socialPB | 0.497241 | 0.690728 | NA | 1.083104 |
| | sexF:socialSol | 0.048999 | 0.364969 | 0.114944 | NA |

*Key*: post. mean = posterior mean, l-95 % = lower 95% credible interval limit, u-95 % = upper 95% credible interval limit, pMCMC = Bayesian two-tailed p-value at alpha = 0.05. *Brain region abbreviations:* teleost: telen. = telencephalon, Dm = medial part of the dorsal telen., Vd = dorsal part of the ventral telen., Dl = lateral part of the dorsal telen., Vv/Vl = lateral and ventral part of the ventral telen., Vs = supracommissural part of the ventral telen., Vc = central part of the ventral telen., POA = pre optic area, TPp = periventricular part of the posterior tuberculum., putative mammalian homolog: blAMY = basolateral amygdala, NAcc = nucleus accumbens, HIP = hippocampus, LS = lateral septum, meAMY/BNST = medial amygdala/bed nucleus of the stria terminalis, Str = Striatum, CP = caudate putamen, VTA = ventral tegmental area.

**Supplementary Table S5:** Model summary of gene expression differences between male *Chaetodon* butterflyfish species within the supracommisural part of the ventral telencephalon. Results are reported as natural log-fold changes of posterior mean from the *a priori* comparison state of *Chaetodon baronessa* with a two-tailed p value.

| **Vs (meAMY.BNST)** | post.mean | l-95% | u-95% | eff.samp | pMCMC |
|---|---|---|---|---|---|
| geneD1R | -3.85E+00 | -9.19E+00 | 8.24E-03 | 341.655 | 0.02 |
| geneD2R | 1.33E+00 | -7.12E-01 | 3.00E+00 | 1000 | 0.162 |
| geneITR | -2.78E+00 | -5.25E+00 | -3.77E-01 | 748.042 | 0.006 |
| geneMOR | 3.18E+00 | 1.21E+00 | 5.18E+00 | 1000 | 0.004 |
| geneV1aR | -2.68E+00 | -4.98E+00 | -5.10E-01 | 1000 | 0.008 |
| gener18S | 1.29E+01 | 1.20E+01 | 1.38E+01 | 1000 | <0.001 |
| geneD1R:speciesC.lun | 4.75E+00 | 4.18E-02 | 1.03E+01 | 581.696 | 0.03 |
| geneD2R:speciesC.lun | 1.75E+00 | -1.16E+00 | 4.40E+00 | 1000 | 0.204 |
| geneITR:speciesC.lun | 7.14E-01 | -2.70E+00 | 4.16E+00 | 794.09 | 0.68 |
| geneMOR:speciesC.lun | -4.88E-01 | -4.06E+00 | 2.52E+00 | 1000 | 0.746 |
| geneV1aR:speciesC.lun | 2.47E+00 | -4.68E-01 | 5.64E+00 | 1000 | 0.102 |
| gener18S:speciesC.lun | 3.61E-01 | -6.57E-01 | 1.44E+00 | 1000 | 0.514 |
| geneD1R:speciesC.rainf | 5.44E+00 | -5.06E-01 | 1.23E+01 | 621.632 | 0.042 |
| geneD2R:speciesC.rainf | -8.82E-01 | -4.74E+00 | 2.65E+00 | 980.066 | 0.654 |

| | | | | | |
|---|---|---|---|---|---|
| geneITR:speciesC.rainf | -9.77E+01 | -1.66E+02 | 1.97E+00 | 4.575 | 0.006 |
| geneMOR:speciesC.rainf | -3.03E+00 | -7.46E+00 | 9.41E-01 | 1000 | 0.134 |
| geneV1aR:speciesC.rainf | -1.11E+02 | -2.54E+02 | -1.36E+01 | 3.45 | <0.001 |
| gener18S:speciesC.rainf | -1.48E-01 | -1.33E+00 | 9.30E-01 | 1000 | 0.814 |
| geneD1R:speciesC.trif | 6.98E+00 | 9.17E-01 | 1.37E+01 | 491.553 | 0.008 |
| geneD2R:speciesC.trif | 2.38E+00 | -1.36E+00 | 5.81E+00 | 1000 | 0.182 |
| geneITR:speciesC.trif | -6.03E+01 | -1.29E+02 | -2.44E+00 | 5.321 | 0.01 |
| geneMOR:speciesC.trif | 1.76E-01 | -4.20E+00 | 3.98E+00 | 1000 | 0.916 |
| geneV1aR:speciesC.trif | 1.29E+00 | -3.28E+00 | 5.68E+00 | 1000 | 0.542 |
| gener18S:speciesC.trif | 9.37E-03 | -1.02E+00 | 1.21E+00 | 1000 | 0.98 |
| geneD1R:speciesC.vag | 5.34E+00 | 9.25E-01 | 1.10E+01 | 421.043 | 0.008 |
| geneD2R:speciesC.vag | 2.63E+00 | 3.68E-01 | 5.06E+00 | 1000 | 0.032 |
| geneITR:speciesC.vag | 1.40E+00 | -1.50E+00 | 4.58E+00 | 866.362 | 0.382 |
| geneMOR:speciesC.vag | 2.18E+00 | -5.11E-01 | 4.90E+00 | 1000 | 0.09 |
| geneV1aR:speciesC.vag | 3.83E+00 | 1.17E+00 | 6.74E+00 | 1305.185 | 0.004 |
| gener18S:speciesC.vag | 3.65E-01 | -7.42E-01 | 1.28E+00 | 1000 | 0.482 |

**Vs (meAMY.BNST) Pair-wise p values**

| | | | | | | |
|---|---|---|---|---|---|---|
| $D1R | | | | | | |
| | difference | | | | | |
| pvalue | | C.bar | C.lun | C.rainf | C.trif | C.vag |
| | C.bar | NA | 6.846559 | 7.849495 | 10.06954 | 7.699104 |
| | C.lun | 0.072924 | NA | 1.002936 | 3.22298 | 0.852545 |
| | C.rainf | 0.095073 | 0.808614 | NA | 2.220044 | -0.15039 |
| | C.trif | 0.033875 | 0.438651 | 0.650826 | NA | -2.37043 |
| | C.vag | 0.041933 | 0.779403 | 0.968907 | 0.541053 | NA |
| $D2R | | | | | | |
| | difference | | | | | |
| pvalue | | C.bar | C.lun | C.rainf | C.trif | C.vag |
| | C.bar | NA | 2.528046 | -1.27193 | 3.434479 | 3.791663 |
| | C.lun | 0.208957 | NA | -3.79998 | 0.906433 | 1.263617 |
| | C.rainf | 0.651541 | 0.16869 | NA | 4.706412 | 5.063596 |
| | C.trif | 0.186504 | 0.741509 | 0.165786 | NA | 0.357184 |
| | C.vag | 0.030836 | 0.513748 | 0.059955 | 0.887718 | NA |
| $ITR | | | | | | |
| | difference | | | | | |
| pvalue | | C.bar | C.lun | C.rainf | C.trif | C.vag |
| | C.bar | NA | 1.030598 | -140.947 | -86.9908 | 2.018993 |
| | C.lun | 0.687678 | NA | -141.978 | -88.0214 | 0.988395 |
| | C.rainf | 0.047687 | 0.045785 | NA | 53.95665 | 142.9664 |
| | C.trif | 0.11556 | 0.111104 | 0.601884 | NA | 89.00975 |
| | C.vag | 0.388087 | 0.690695 | 0.044372 | 0.106807 | NA |
| $MOR | | | | | | |
| | difference | | | | | |

| pvalue | | C.bar | C.lun | C.rainf | C.trif | C.vag |
|---|---|---|---|---|---|---|
| | C.bar | NA | -0.70369 | -4.37093 | 0.25317 | 3.142817 |
| | C.lun | 0.767663 | NA | -3.66724 | 0.95686 | 3.846507 |
| | C.rainf | 0.159047 | 0.265396 | NA | 4.624097 | 7.513744 |
| | C.trif | 0.933398 | 0.7739 | 0.235666 | NA | 2.889647 |
| | C.vag | 0.106342 | 0.099076 | 0.013392 | 0.338936 | NA |
| $V1aR | | | | | | |
| | difference | | | | | |
| pvalue | | C.bar | C.lun | C.rainf | C.trif | C.vag |
| | C.bar | NA | 3.559984 | -160.411 | 1.856114 | 5.531158 |
| | C.lun | 0.114231 | NA | -163.971 | -1.70387 | 1.971174 |
| | C.rainf | 0.081732 | 0.0756 | NA | 162.2671 | 165.9422 |
| | C.trif | 0.562972 | 0.59314 | 0.078831 | NA | 3.675044 |
| | C.vag | 0.007693 | 0.327244 | 0.072071 | 0.21083 | NA |
| $r18S | | | | | | |
| | difference | | | | | |
| pvalue | | C.bar | C.lun | C.rainf | C.trif | C.vag |
| | C.bar | NA | 0.520708 | -0.21374 | 0.013524 | 0.52661 |
| | C.lun | 0.508665 | NA | -0.73445 | -0.50718 | 0.005902 |
| | C.rainf | 0.796529 | 0.525246 | NA | 0.227265 | 0.740351 |
| | C.trif | 0.987061 | 0.653522 | 0.847257 | NA | 0.513086 |
| | C.vag | 0.47746 | 0.995432 | 0.499484 | 0.638388 | NA |

*Key*: post. mean = posterior mean, l-95 % = lower 95% credible interval limit, u-95 % = upper 95% credible interval limit, pMCMC = Bayesian two-tailed p-value at alpha = 0.05, *Brain region abbreviations:* Vs = supracommissural part of the ventral telencephalon, meAMY/BNST = medial amygdala/bed nucleus of the stria terminalis.
